# Leptomeningeal Enhancement in Multiple Sclerosis and Other Neurological Diseases: A Systematic Review and Meta-Analysis

**DOI:** 10.1101/2021.12.15.472797

**Authors:** Benjamin V. Ineichen, Charidimos Tsagkas, Martina Absinta, Daniel S. Reich

**Affiliations:** Translational Neuroradiology Section, National Institute of Neurological Disorders and Stroke, National Institutes of Health, Bethesda, MD, 20892, United States; Department of Neuroradiology, Clinical Neuroscience Center, University Hospital Zurich, University of Zurich, Switzerland; Neurologic Clinic and Policlinic, Departments of Medicine, Clinical Research and Biomedical Engineering, University Hospital Basel and University of Basel, Basel, Switzerland; Translational Imaging in Neurology (ThINk) Basel, Department of Medicine and Biomedical Engineering, University Hospital Basel and University of Basel, Basel, Switzerland; Department of Neurology, Johns Hopkins University School of Medicine, Baltimore, MD, USA; Vita-Salute San Raffaele University, and Neurology Unit, IRCCS San Raffaele Scientific Institute, Milan, Italy

**Keywords:** multiple sclerosis, leptomeningeal enhancement, leptomeningeal inflammation, magnetic resonance imaging, systematic review, meta-analysis

## Abstract

**Background:** The lack of systematic evidence on leptomeningeal enhancement (LME) on MRI in neurological diseases, including multiple sclerosis (MS), hampers its interpretation in clinical routine and research settings.

**Purpose:** To perform a systematic review and meta-analysis of MRI LME in MS and other neurological diseases.

**Materials and Methods:** In a comprehensive literature search in Medline, Scopus, and Embase, out of 2292 publications, 459 records assessing LME in neurological diseases were eligible for qualitative synthesis. Of these, 135 were included in a random-effects model meta-analysis with subgroup analyses for MS.

**Results:** Of eligible publications, 161 investigated LME in neoplastic neurological (n=2392), 91 in neuroinfectious (n=1890), and 75 in primary neuroinflammatory diseases (n=4038). The LME-proportions for these disease classes were 0.47 [95%-CI: 0.37–0.57], 0.59 [95%-CI: 0.47–0.69], and 0.26 [95%-CI: 0.20–0.35], respectively. In a subgroup analysis comprising 1605 MS cases, LME proportion was 0.30 [95%-CI 0.21–0.42] with lower proportions in relapsing-remitting (0.19 [95%-CI 0.13–0.27]) compared to progressive MS (0.39 [95%-CI 0.30–0.49], p=0.002) and higher proportions in studies imaging at 7T (0.79 [95%-CI 0.64–0.89]) compared to lower field strengths (0.21 [95%-CI 0.15–0.29], p<0.001). LME in MS was associated with longer disease duration (mean difference 2.2 years [95%-CI 0.2–4.2], p=0.03), higher Expanded Disability Status Scale (mean difference 0.6 points [95%-CI 0.2–1.0], p=0.006), higher T1 (mean difference 1.6ml [95%-CI 0.1–3.0], p=0.04) and T2 lesion load (mean difference 5.9ml [95%-CI 3.2–8.6], p<0.001), and lower cortical volume (mean difference −21.3ml [95%-CI −34.7–-7.9], p=0.002).

**Conclusions:** Our study provides high-grade evidence for the substantial presence of LME in MS and a comprehensive panel of other neurological diseases. Our data could facilitate differential diagnosis of LME in clinical settings. Additionally, our meta-analysis corroborates that LME is associated with key clinical and imaging features of MS.

PROSPERO No: CRD42021235026.

**Summary statement:** Our systematic review and meta-analysis synthesize leptomeningeal enhancement proportions across a comprehensive panel of neurological diseases, including multiple sclerosis, and assesses its prognostic value in multiple sclerosis.

**Summary data:**

- Leptomeningeal enhancement (LME) is a nonspecific imaging feature present across many neurological disorders, including neoplasm, infection, and primary neuroinflammation.
- The presence of LME is associated with worse clinical and imaging outcomes in multiple sclerosis, justifying its ascertainment in clinical practice.
- Neuroinflammatory animal models can be used to further investigate the pathophysiology of LME, including its pathological tissue signature and/or its association with cortical pathology.

## Introduction

Abnormal meningeal contrast enhancement may take two distinct forms: pachymeningeal enhancement, referring to dural-arachnoidal enhancement, which follows the contour of the inner table of the skull and includes intradural veins and sinuses; and leptomeningeal enhancement (LME), which follows the pia-arachnoid abutting the cortical surface and extending into the sulci. LME is often caused by neoplastic or infectious processes. However, LME is also gaining increased attention as a putative imaging biomarker of meningeal inflammation in neuroinflammatory diseases, including MS and neurosarcoidosis (**Figure 1**).(1)

**Figure 1:**
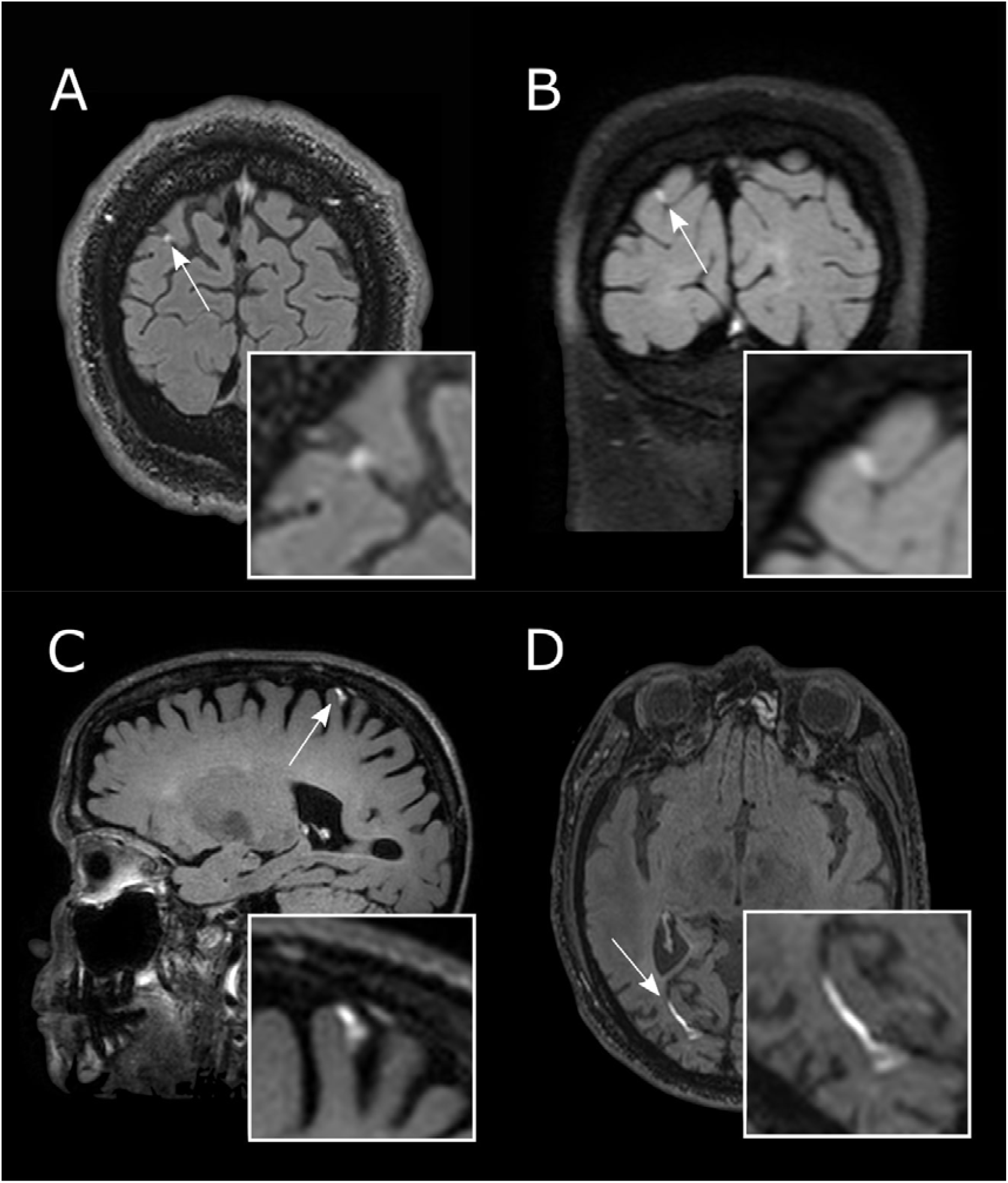
Leptomeningeal enhancement (LME) across the spectrum of neurological diseases. Leptomeningeal enhancement (LME, white arrows) can be detected using post-gadolinium fluid-attenuated inversion recovery (FLAIR) T2-weighted magnetic resonance imaging and can be present in viral diseases such as HIV (A, at 3T), in primary neuroinflammatory diseases such as Susac syndrome (B, at 3T) and multiple sclerosis (C, at 7T), and in aseptic meningitis (natalizumab-induced) (D, at 3T).

Because LME can be present in a wide variety of neurological diseases,(2) differential diagnostic considerations are paramount for proper patient workup. However, there is a lack of systematic evidence on LME proportions in MS and other neurological diseases. Furthermore, a wide variety of methodological approaches to imaging LME has been published,(3, 4) impeding the implementation of an appropriate imaging protocol for sensitive LME detection. With respect to MS, several studies have presented conflicting findings regarding the association of LME with clinical and imaging parameters,(5) such that high-level evidence would benefit clinicians and researchers.

Based on these shortcomings, we set out to systematically summarize the available evidence on LME in neurological diseases with a focus on MS. This study had the following goals: (1) synthesize data on LME proportions in neurological diseases, including potentially distinct LME features such as phenotype or temporal evolution; (2) qualitatively and quantitatively summarize the potential association of LME with clinical and imaging features in MS; (3) propose an appropriate imaging protocol to detect LME in clinical and research practice; (4) summarize the data on pathological correlates of LME in neuroinflammation; (5) summarize the available evidence on LME in animal models of neuroinflammation.

## Materials and Methods

We registered the study protocol in the International prospective register of systematic reviews (PROSPERO, CRD42021235026, https://www.crd.york.ac.uk/PROSPERO/) and used the Preferred Reporting Items for Systematic Reviews and Meta-Analysis (PRISMA) Guidelines for reporting.(6)

### Search strategy

We searched for original studies published in full up to February 2, 2021, in PubMed, Scopus, and Ovid EMBASE. See **Table S1** for the search string in each of these databases.

### Inclusion and exclusion criteria

We included publications on human or animal data that reported on any outcome related to leptomeningeal inflammation on magnetic resonance imaging (MRI) in any neurological diseases. Case reports were also included in the systematic review. Exclusion criteria: conference abstracts, non-English articles, and publications that reiterated previously reported quantitative data. Reviews were excluded but retained as potential sources of additional records.

### Study selection and data extraction

Titles and abstracts of studies were screened for their relevance in the web-based application Rayyan by two reviewers (CT and BVI), followed by full-text screening.(7) Subsequently, the following data were extracted: title, authors, publication year, study design, neurological disease, and number of subjects per group. For studies with ≥10 subjects, MRI sequences/field strength, LME location (spinal cord, convexities, basal, cerebellar, brainstem), main study findings (in narrative manner), pattern of LME (for example, nodular or linear), temporal dynamics of LME, and proportions of LME in experimental and control groups were also extracted.

### Quality assessment

The quality of each study with ≥10 included subjects was assessed against predefined criteria by two reviewers (CT and BVI) using the Newcastle-Ottawa scale for evaluating risk of bias in nonrandomized studies.(8) Discrepancies were resolved by discussion.

### Data synthesis and analysis

Only diseases/disease classes with ≥2 publications describing ≥10 adult subjects each were included in the meta-analysis, and only summary-level data were used. As primary outcome, log-transformed proportions of LME were used. A random-effects model was fitted to the data. The amount of heterogeneity (τ2), was estimated using the DerSimonian-Laird estimator.(9) In addition, the Q-test for heterogeneity (10) and the I^2^ statistic (11) are reported.

For subgroup analyses of clinical and imaging outcomes in MS, the analysis was carried out using the log-transformed proportions or mean difference as the outcome measure. Subgroup analyses were computed for MS to assess the proportion of LME in clinical MS phenotypes when ≥3 studies were available. Subgroup analyses in MS for clinical and imaging outcomes were computed when ≥3 studies reported at least mean, variance, and n for LME+ and LME-groups on respective outcomes.

A two-tailed P value <0.05 was considered statistically significant.

### Publication bias

The rank correlation test and the regression test, using the standard error of the observed outcomes as predictor, were used to check for funnel plot asymmetry. The analysis was carried out using R (version 3.6.1) with the *meta* and *metafor* packages (version 2.4.0).(12)

## Results

### 1. Eligible publications

In total, 2292 original publications were retrieved from our comprehensive database search and an additional 10 publications from reference lists of reviews on related topics. After abstract and title screening, 1089 studies were eligible for full-text search. After screening the full text of these studies, 458 articles (35% of deduplicated references) were included for qualitative synthesis and 135 articles (10%) for quantitative synthesis (**Figure S1**).

### 2. General study characteristics

#### 2.1. Included publications

Of the eligible publications, 144 investigated LME in neoplastic neurological diseases (2392 subjects including 183 children), 91 in infectious neurological diseases (1890 subjects including 48 children), and 76 in primary neuroinflammatory diseases (4038 subjects including 11 children). Additionally, 147 publications assessed LME in neurological diseases that did not belong to mentioned categories (1961 subjects including 762 children). We also included 5 publications in animal models of neurological diseases (mouse experimental autoimmune encephalomyelitis [EAE] model, bacterial meningitis in rats, bacterial CNS infection in a dog, subarachnoid diverticulum in a cat).

#### 2.2. Risk of bias assessment

Most studies showed a low risk of bias for the selection domain (that is, whether patients and controls were defined according to acknowledged diagnostic criteria); see **Table S2 and S3.** Many studies did not report on adjusting their statistical analyses for subject age, sex, or other potential confounders (comparability domain), thus potentially inducing biases.

### 3. Leptomeningeal enhancement in neuroinflammatory diseases including multiple sclerosis

#### 3.1. Primary neuroinflammatory diseases overall

##### 3.1.1. Diseases

Studies reporting on LME in neuroinflammatory diseases were in neurosarcoidosis (20 publications), MS (17 publications), MOG-antibody diseases/encephalitis (8 publications), neuromyelitis optica spectrum disorder (NMOSD) (8 publications), primary angiitis of the CNS (6 publications), Susac syndrome (5 publications), NMDA-receptor encephalitis (4 publications), Behçet syndrome (1 publication), and GFAP astrocytopathy (1 publication).

##### 3.1.2. LME pattern

21 studies did not report on the LME pattern, whereas the remaining studies reported on different LME patterns. The pattern of LME has most extensively been described in MS in 12 publications, mostly as either nodular and/or laminar/spread-and-fill.(13–19) Similar LME patterns have been described in Susac syndrome (20) and neurosarcoidosis.(21, 22) A spreading/laminar phenotype has been described in NMOSD,(23–25) NMDA-receptor encephalitis,(26) and GFAP astrocytopathy.(27)

##### 3.1.3. MRI acquisition of LME

Most studies employed a postcontrast T2w-FLAIR sequence to visualize LME (19 publications) followed by a postcontrast T1w sequence (6 publications). 11 studies did not report the sequences used to detect LME.

##### 3.1.4. Meta-analysis

A meta-analysis of LME in primary neuroinflammatory diseases, including a total of 2284 patients, showed an overall proportion of 0.26 [95%-CI: 0.20–0.35] with substantial heterogeneity across studies (I^2^=90%, p<0.001) (**Figure 2**). NMOSD had the lowest proportion of LME with 0.06 [95%-CI 0.02–0.23], and neurosarcoidosis had the highest LME proportion with 0.41 [95%-CI 0.13–0.52].

**Figure 2:**
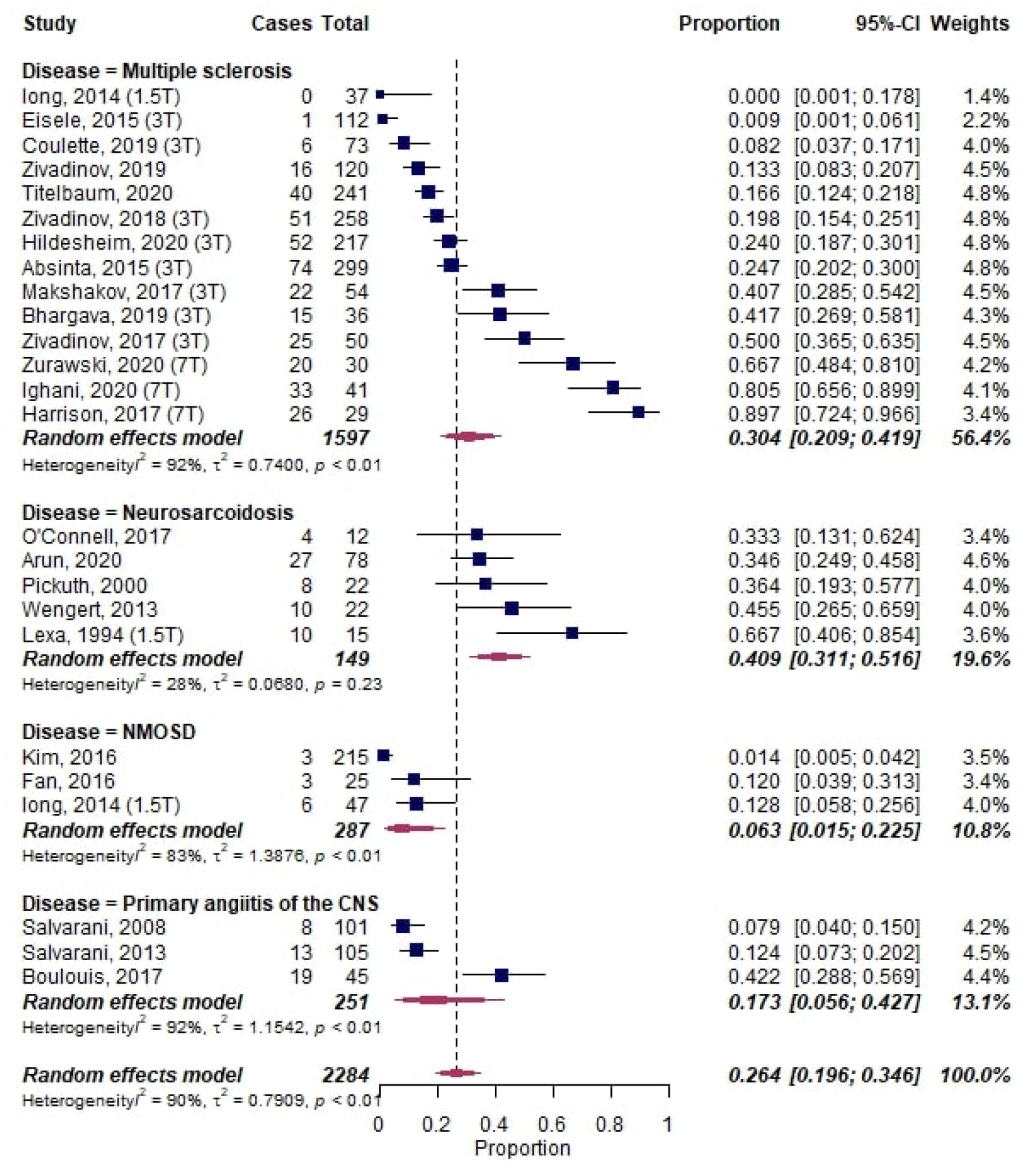
Forest plot of leptomeningeal enhancement (LME) proportions in primary neuroinflammatory diseases. Pooled analyses of studies comparing the proportion of LME on MRI in neuroinflammatory diseases, stratified by diseases. Static magnetic field strength for MRI acquisition for respective studies are listed in brackets if reported. Proportions for LME were extracted and pooled using the random effects DerSimonian-Laird method. Abbreviations: CNS, central nervous system; CI, confidence interval; NMOSD, neuromyelitis optica spectrum disorder.

#### 3.2. Multiple sclerosis

##### 3.2.1. LME proportion in MS and subgroups

Two studies from 2015 first described LME in MS.(2, 28) These and subsequent studies comprised 1605 MS patients, 303 non-MS controls, and 126 healthy controls (**Table 1**).

**Table 1:**
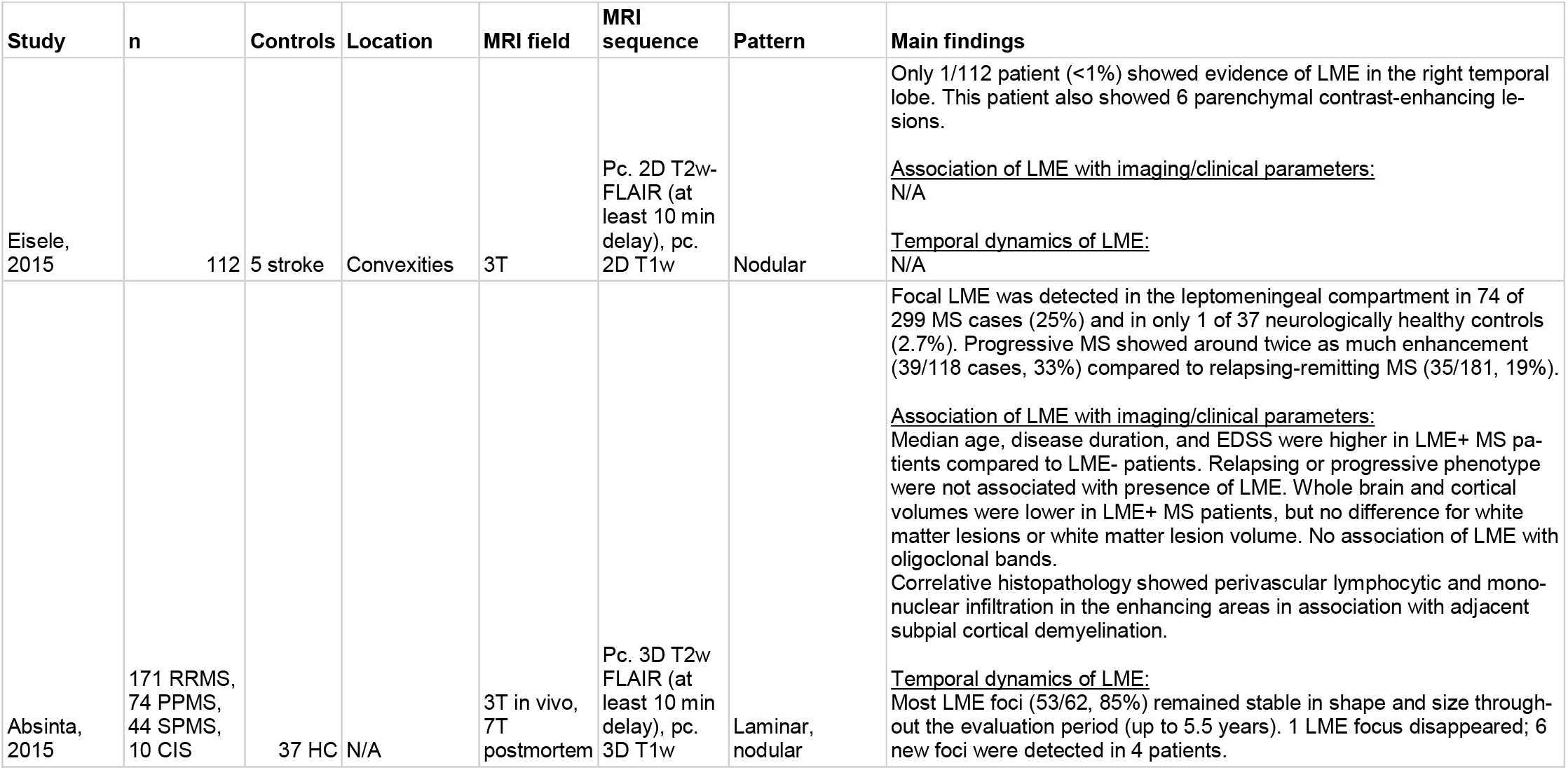
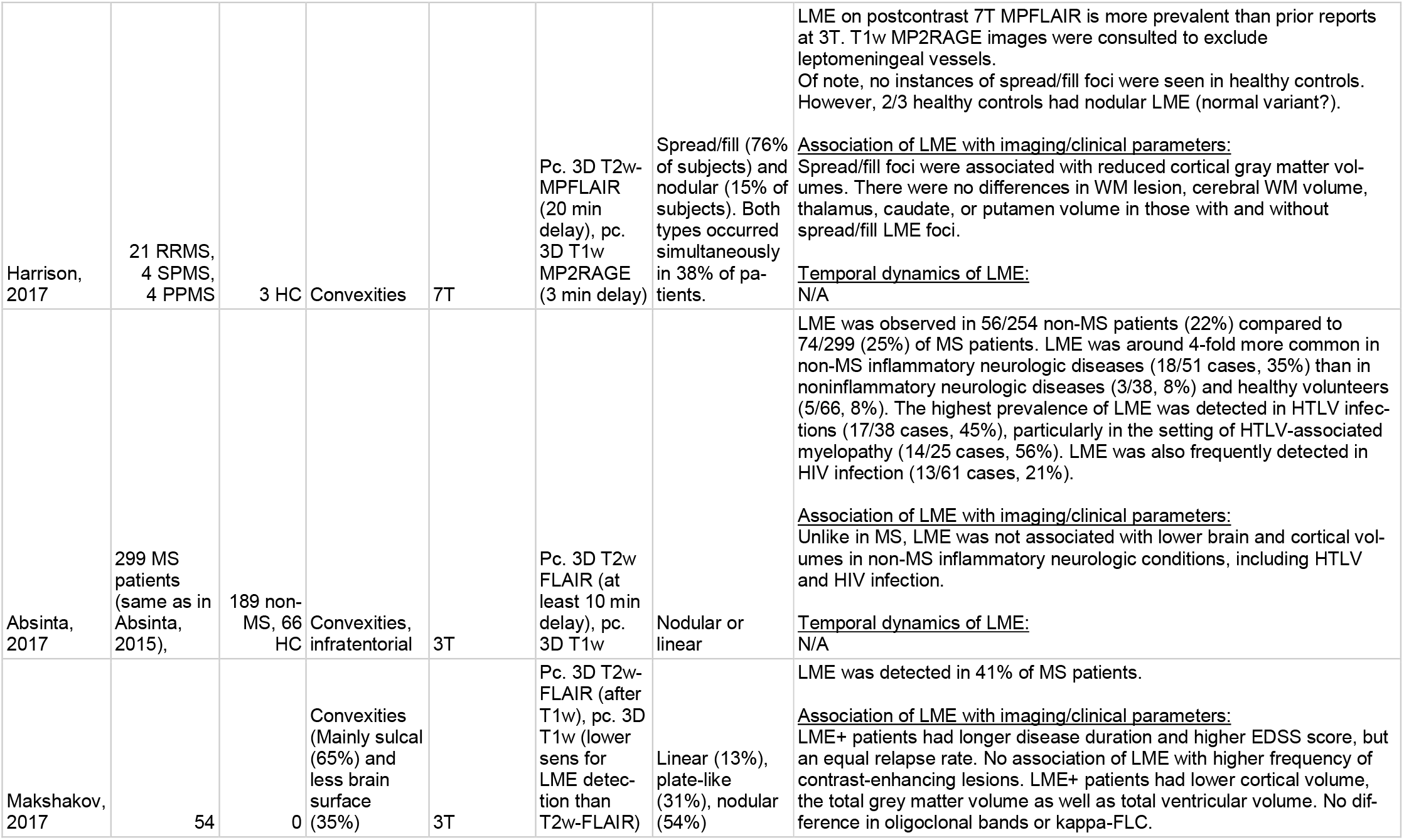
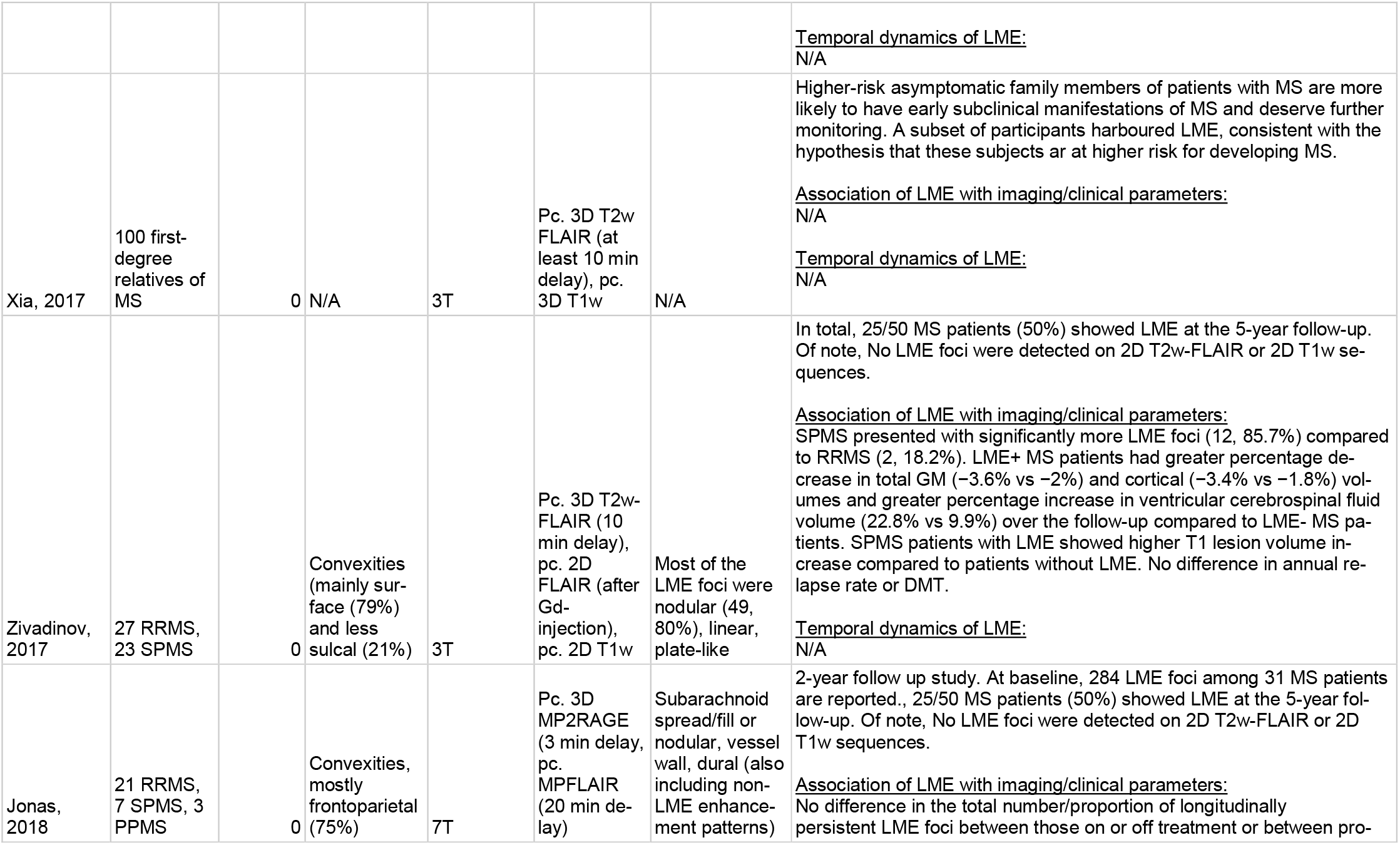
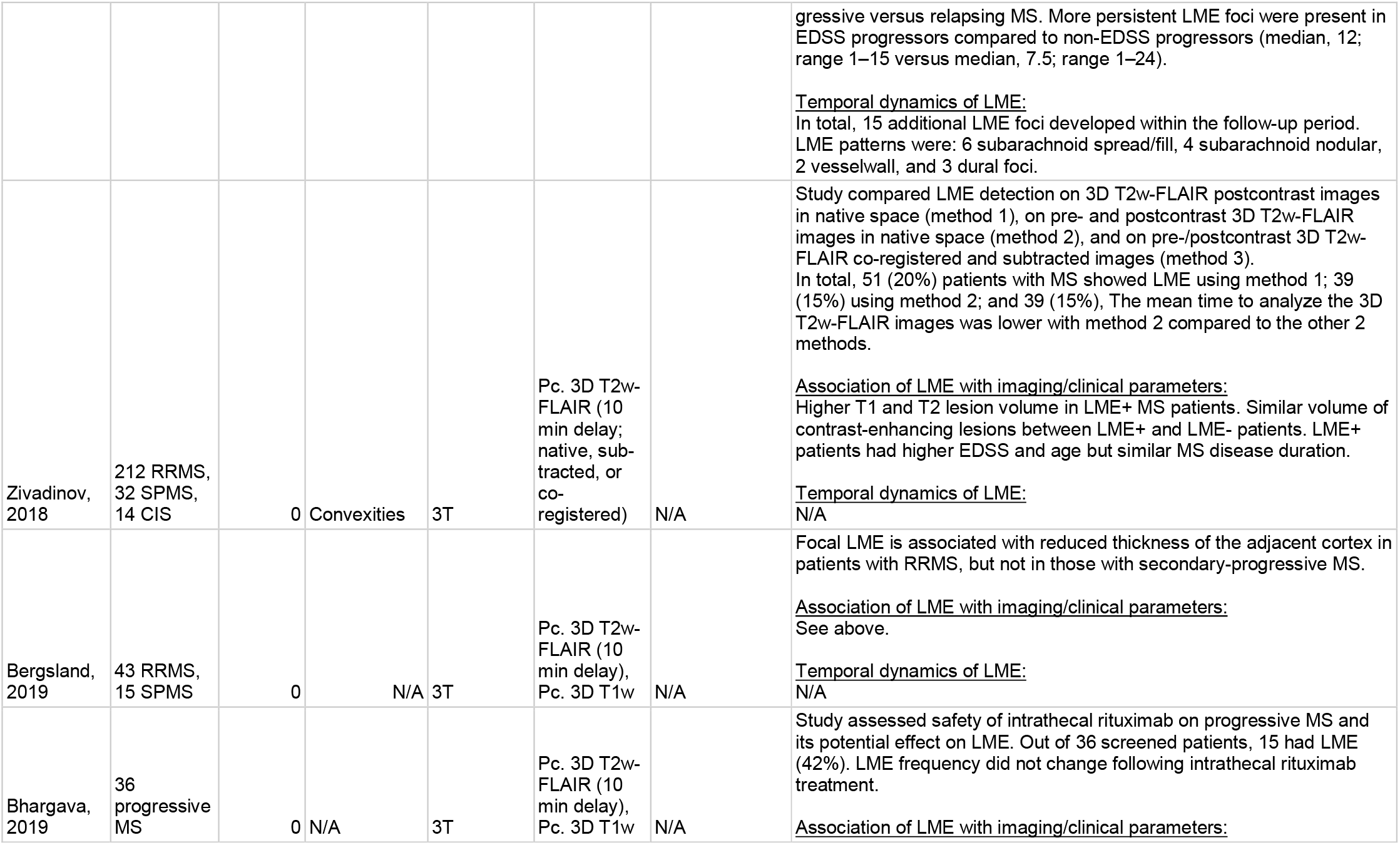
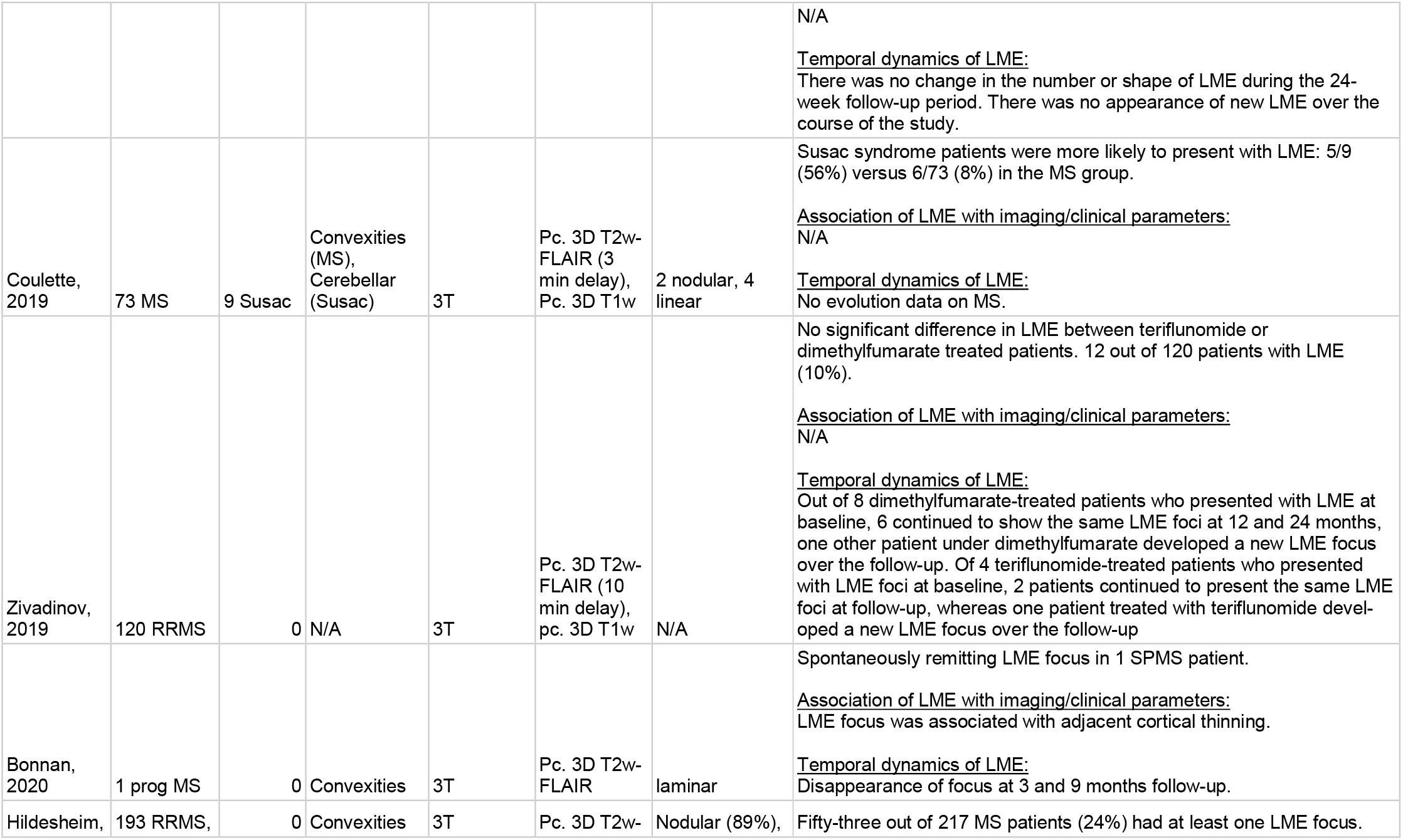
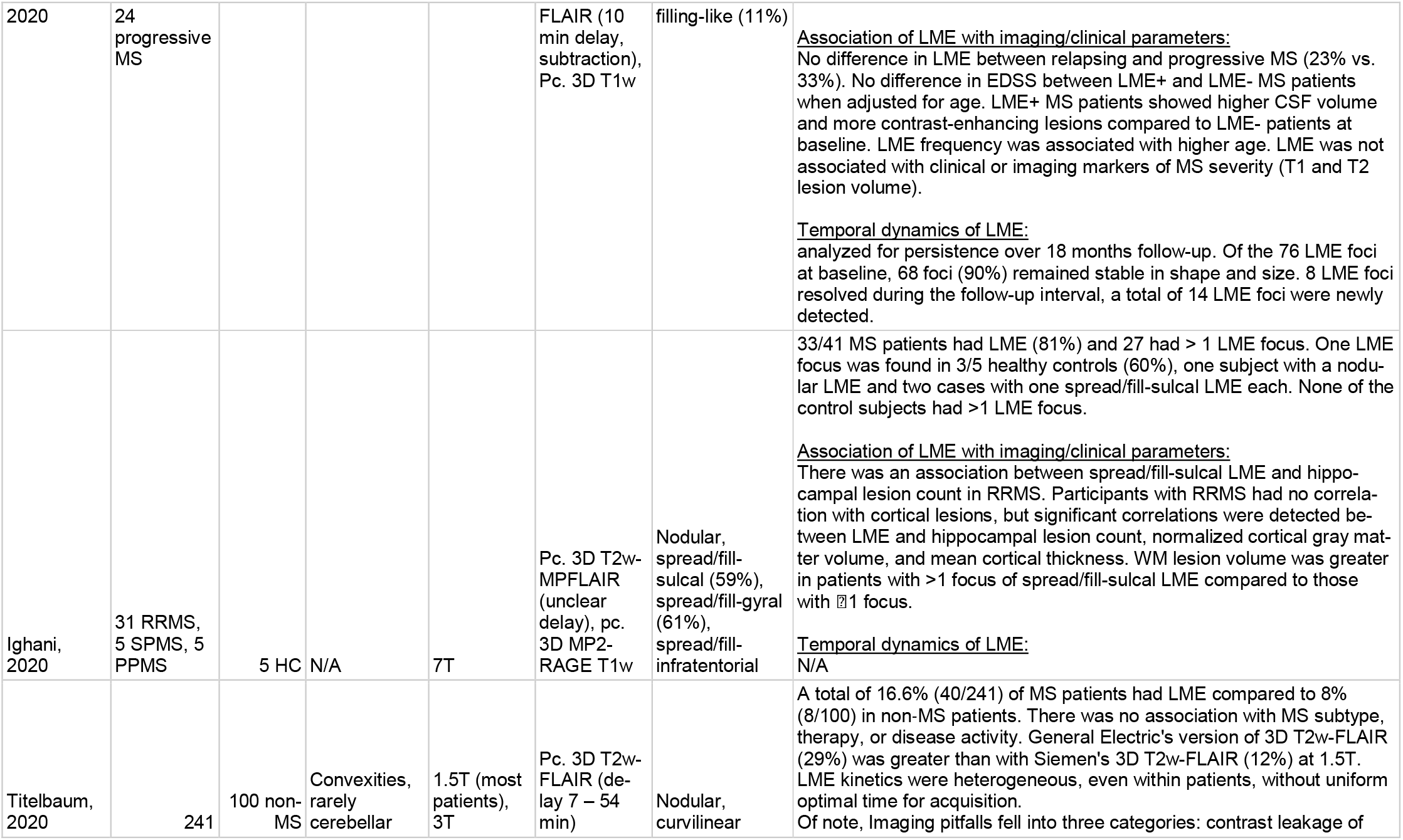
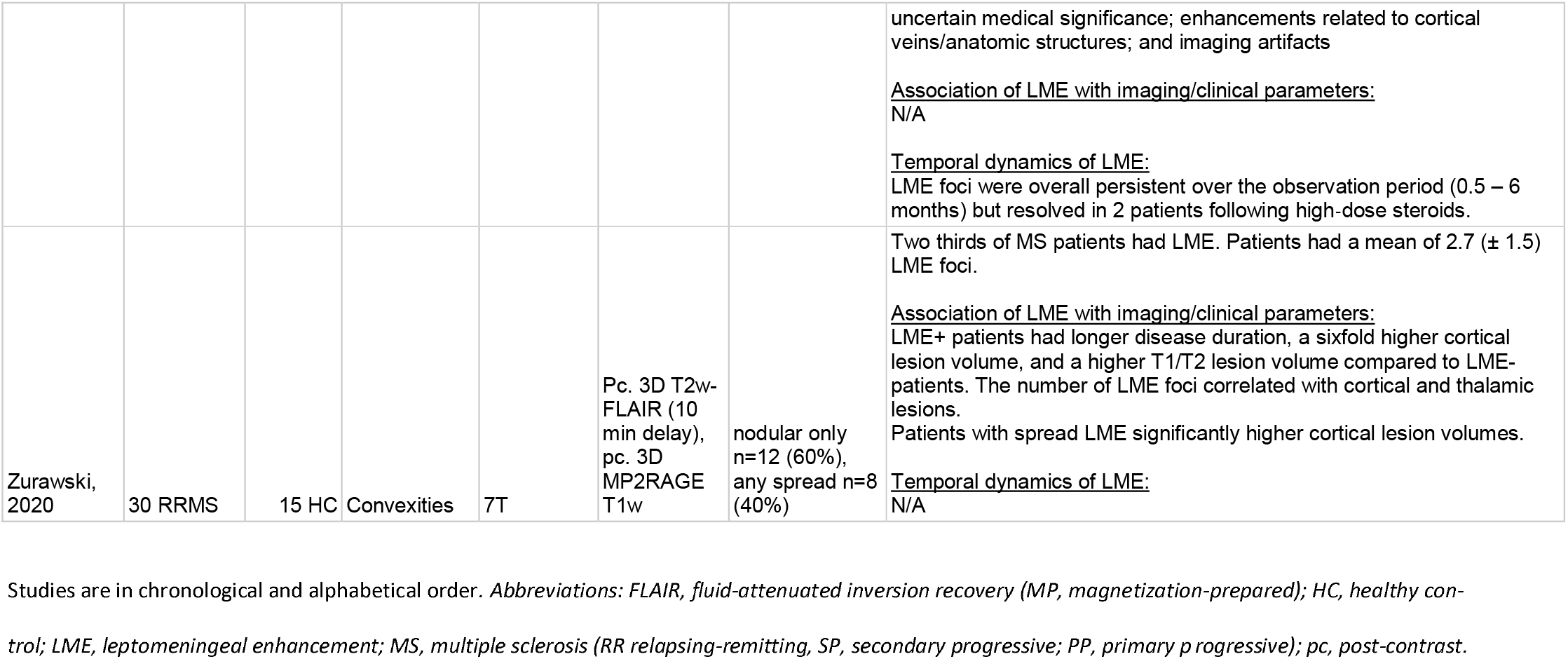
Synopsis of studies assessing leptomeningeal enhancement (LME) in multiple sclerosis (MS).

The overall proportion of LME in MS was 0.30 [95%-CI 0.21–0.42] (**Figure 3**). However, LME proportions in MS patients with a relapsing-remitting clinical phenotype (0.19 [95%-CI 0.13–0.27]; 7 publications) and CIS patients (0.06 [95%-CI 0.02–0.20]; 3 publications) were significantly lower compared to progressive MS patients (0.39 [95%-CI 0.30–0.49], p=0.002 and p=0.003, respectively; 6 publications).

**Figure 3:**
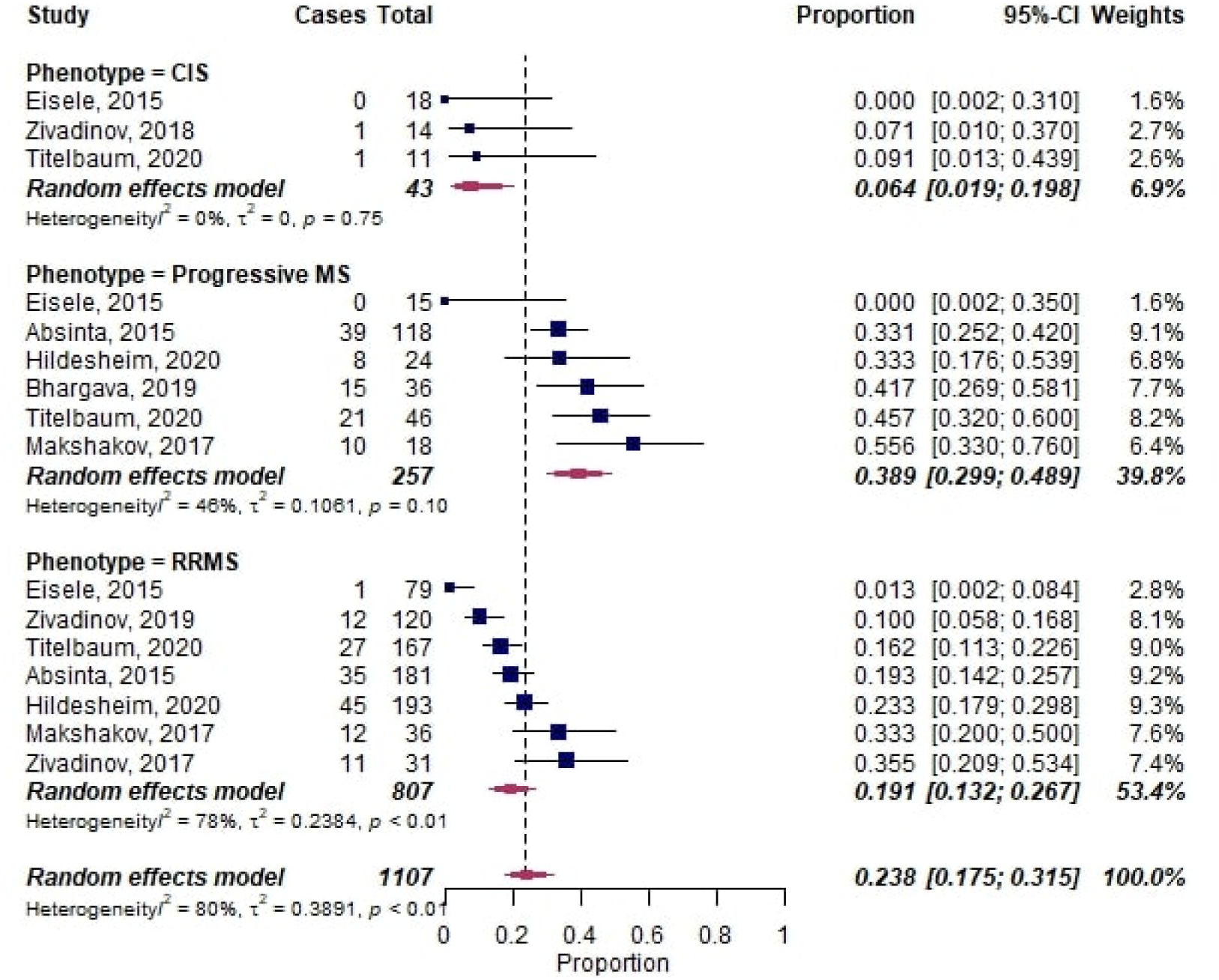
Forest plot of leptomeningeal enhancement (LME) proportions in multiple sclerosis (MS). Pooled analyses of studies comparing the proportion of LME on MRI in MS, stratified by clinical phenotype. Proportions for LME were extracted and pooled using the random effects DerSimonian-Laird method. Abbreviations: *CI* confidence interval; CIS, clinically isolated syndrome; RRMS, relapsingremitting multiple sclerosis.

##### 3.2.2. MRI acquisition of LME

14 studies acquired MRI at 3T (or complementing 1.5T) and 3 studies at 7T.(13, 16, 18) In a meta subgroup-analysis with the B_0_ magnetic field strength as moderator, LME proportions were higher in studies imaging at 7T (0.79 [95%-CI 0.64–0.89]; 3 publications) compared to studies imaging at 1.5/3T (0.21 [95%-CI 0.15–0.29], p<0.001; 11 publications) (**Figure 4**)

**Figure 4:**
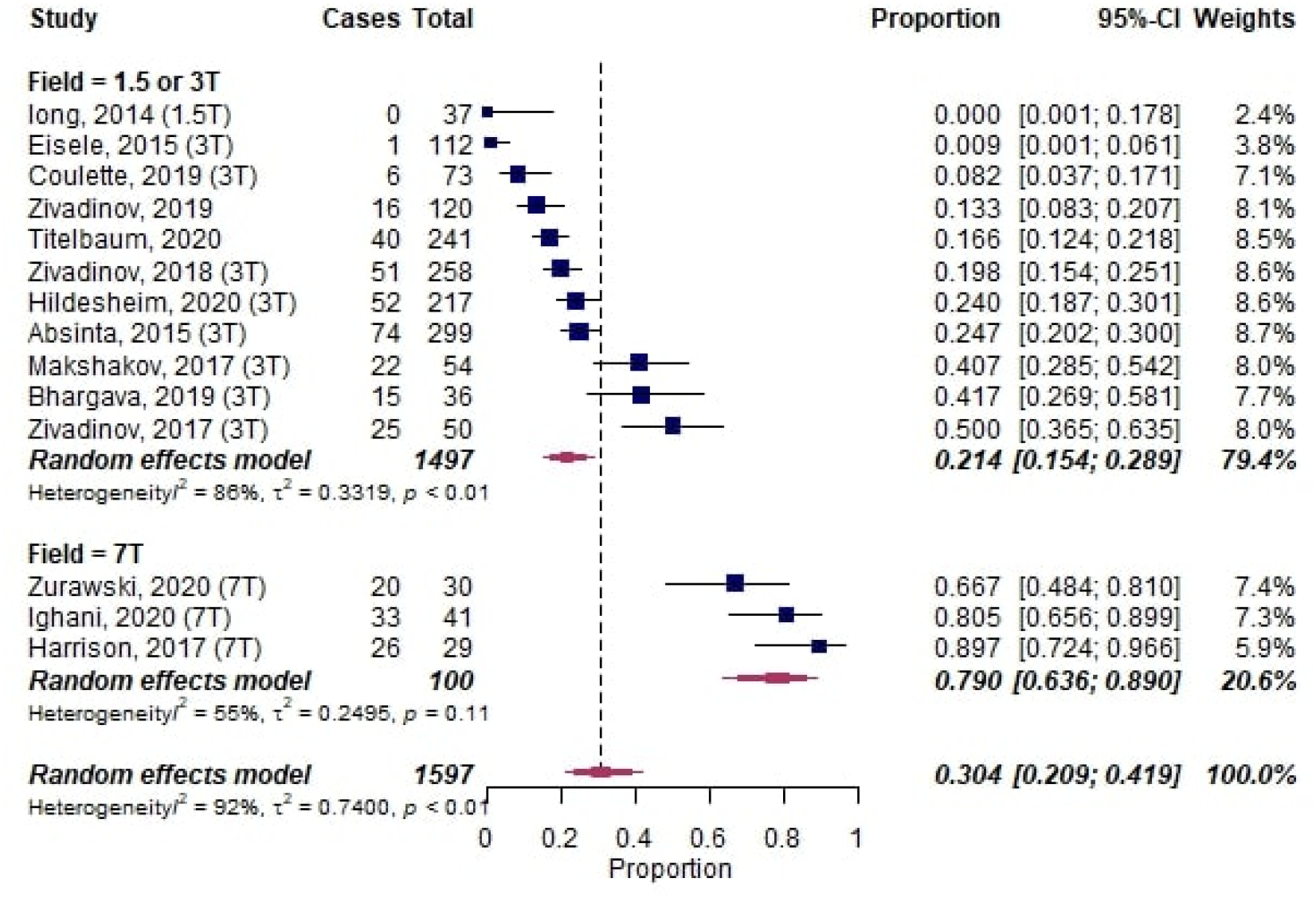
Forest plot of leptomeningeal enhancement (LME) proportions in multiple sclerosis (MS) with B_0_ magnetic field strength as moderator. Pooled analyses of studies comparing the proportion of LME on MRI in MS, stratified by magnetic field strength. Proportions for LME were extracted and pooled using the random effects DerSimonian-Laird method. Abbreviations: *CI* confidence interval.

All 17 studies investigating LME in MS employed both a postcontrast 3D T2w-FLAIR and a postcontrast 3D T1w sequence to detect LME. Of note, 1 study acquired 2D scans instead of 3D, showing the lowest LME proportion among all MS studies (< 0.01, 1/112 cases).(28) Of note, is has been shown that 3D T2w-FLAIR can have higher sensitivity to detect gadolinium enhancement compared to T1w imaging, particularly for superficial enhancement.(29) The acquisition of the 3D T2w-FLAIR sequence was mostly 10 minutes after Gd-injection, with two studies reporting either earlier (3 minutes)(20) or later acquisition (up to 54 minutes).(15) It is noteworthy that latter study suggested that individual LME foci might have different enhancement kinetics and thus different peak enhancement time points, and that differences in LME detection between scanner types could be due to differences in 3D T2w-FLAIR sequences.

One study found lower sensitivity for LME detection using T1w sequences compared to T2w-FLAIR.(19) One study compared LME detection on 3D T2w-FLAIR postcontrast images in native space (method 1), on pre- and postcontrast 3D T2w-FLAIR images in native space (method 2), and on pre-/postcontrast 3D T2w-FLAIR co-registered and subtracted images (method 3).(3) In total, 51 (20%) MS cases showed LME using method 1; 39 (15%) using method 2; and 39 (15%) using method 3. The mean time to analyze the 3D T2w-FLAIR images was lower with method 2 compared to the other 2 methods.

##### 3.2.3. LME phenotypes

Overall, two configurations of LME have been described, albeit with inhomogeneous nomenclature: nodular and “spread-fill” (subsuming linear, laminar, and plate-like). Most studies described these two phenotypes (12/17 publications).(2, 13-20, 30) 1 early publication with a very low LME proportion of <0.01 exclusively described a nodular LME phenotype.(28) 5 publications did not declare the LME pattern.

The prevalence of these patterns varied considerably between studies: 4 publications observed higher frequencies of nodular LME: 60% (vs. spread 40%),(18) 54% (vs. 13% linear and 31% plate-like),(19) 80% (vs. linear and plate-like),(17) and 89% (vs. 11% filling-like).(14) Two publications described higher frequencies of linear/spread-fill LME: 59–61% for spread-fill sulcal or gyral (vs. nodular)(13) and 76% spread-fill (vs. 15% nodular).(16) The latter study observed simultaneous presence of both LME phenotypes in 38% of MS cases.

##### 3.2.4. Temporal evolution of LME

Eleven studies did not include longitudinal MRI data and did thus not assess temporal evolution of LME. The remaining 6 studies consistently reported mostly stable LME foci over several years. One study with a follow-up period of up to 5.5 years reported that 85% of LME foci remained stable and that only 6 new LME foci were detected over this observation period (in 4 of 299 MS patients).(30) Similar high percentages of stable LME foci have been observed in other studies: 75% stable over 24 months (and 2 new LME foci in 2/120 patients),(31) 90% stable over 18 months (and 14 new LME foci),(14) 100% stable over 24 weeks (32), and 73–100% stable over 24 months (33). The last study also included non-LME enhancement patterns and found that subarachnoid nodular and spread/fill LME patterns persisted less often than dural or vessel wall foci, and that MS patients with EDSS progression showed more persistent LME foci.

##### 3.2.6. Association of LME with clinical and imaging parameters

Three studies found an association between LME and age and/or disease duration (14, 19, 30), which was not confirmed for disease duration by 1 study.(3) An association between LME and Expanded Disability Status Scale (EDSS) was described in 3 studies (3, 19, 30) but was not confirmed in 1 study after adjusting for age.(14) MS relapse rate was not associated with LME in 2 studies.(17, 19)

Five studies assessed the association between LME and T1 or T2 lesion volume. Three studies found such an association,(3, 17, 18) whereas 2 did not.(14, 16) Six studies consistently reported lower cortical gray matter volume and/or thickness in MS cases with LME.(13, 16–19, 34) Two studies assessed the association of LME with cortical MS lesions at 7T; one of these studies found no such association (13) while the other did.(18) Both studies found an association of LME with subcortical gray matter MS lesions (thalamic and hippocampal, respectively).

In view of the inconsistency regarding the association of LME with clinical and imaging parameters in MS, we assessed these effects in a meta-analysis (for all outcomes reported in ≥3 publications). In this analysis, the presence of LME was associated with longer disease duration (mean difference 2.2 years [95%-CI 0.2–4.2], p=0.03, **Figure 5A**) and higher EDSS (mean difference 0.6 EDSS points [95%-CI 0.2–1.0], p=0.006, **Figure 5B**). Additionally, MS cases with LME showed higher T1 lesion volume (mean difference 1.5 ml [95%-CI 0.1–3.0], p=0.04, **Figure 5C**), higher T2 lesion volume (mean difference 5.9 ml [95%-CI 3.2–8.6], p<0.001, **Figure 5D**), and lower cortical volume (mean difference −21.3 ml [95%-CI -−34.7–-7.89], p=0.002, **Figure 5E**).

**Figure 5:**
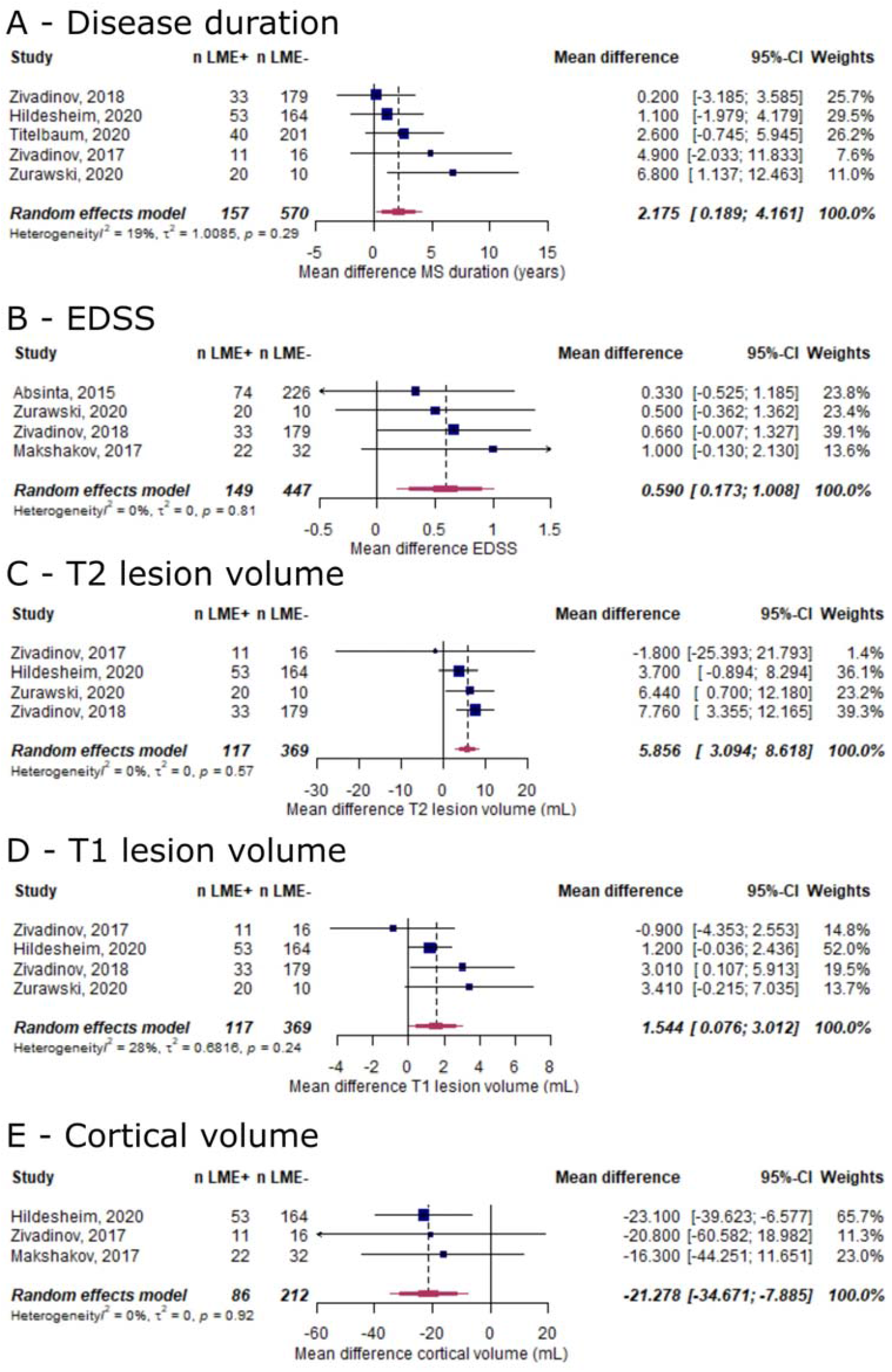
Forest plot for subgroup analysis for the association between LME and clinical and imaging outcomes in MS. Pooled analyses of studies comparing the mean differences of clinical and imaging outcomes between LME-positive and LME-negative MS cases (A, disease duration [years]; B, EDSS [in EDSS points]; C, T1 lesion volume [in mL]; D, T2 lesion volume [in mL]; E, cortical gray matter volume [in mL]). Mean differences were extracted and pooled using the random effects DerSimonian-Laird method. Abbreviations: CI confidence interval; EDSS, Expanded Disability Status Scale; LME, leptomeningeal enhancement.

##### 3.2.7. Histopathological validation of LME

One study performed histopathological validation of three LME foci in two progressive MS cases.(30) The gyri adjacent to the LME foci were affected by confluent cortical demyelination and/or subpial cortical demyelination. Leptomeningeal perivascular inflammation, including T cells, B cells, and macrophages, was detected in these areas.

##### 3.2.7. LME in MS animal models

We included two studies that assessed LME in mouse EAE using postcontrast T2w-FLAIR. The first study employing 9.4T MRI found that all 13 inoculated mice showed LME foci, compared to none of the control mice.(35) Peak LME intensity was at 10 days post induction and correlated with weight loss and clinical symptoms. In a histopathological analysis, LME foci were associated with high densitiy of Iba-1 positive microglia cells as well as T and B cells, which were absent in control mice. Another recent study also emplyoing mouse EAE and MRI at 11.7T with pathological correlation showed that mice treated with a Bruton tyrosine kinase-inhibitor (evobrutinib) had a reduced number of LME foci, while anti-CD20 therapy had no effect on LME.(36) The pathological tissue substrate showed that this corresponded to a reduction in B cells within regions of meningeal inflammation as well as reduced astrocytosis in the adjacent cortex. Interestingly, myeloid cell infiltrates seemed to persist despite B cell depletion.

##### 3.2.5. Therapeutic impact on LME

Four studies assessed the impact of drug treatment on LME resolution. One study found similar LME persistence rates in MS patients with/without disease-modifying therapy (DMT).(33) Another study assessing the efficacy of dimethyl fumarate or teriflunomide on LME reported no differences in LME resolution between treatment groups (8 of 12 patients showed stable LME, and 2 patients developed new LME).(31) One study assessing intrathecal rituximab treatment in 15 progressive MS patients observed stable LME foci in all patients over the 24-week follow-up period.(32) In contrast to these studies with relatively small sample size and consequently lower statistical power, one study including 241 MS patients observed resolution of LME in 2 patients after high-dose steroid treatment within 6 months follow-up.(15)

### 4. Leptomeningeal enhancement in other neurological diseases

#### 4.1. Infectious CNS diseases

##### 4.1.1. Diseases

Studies reported on LME in infectious (encephalo-)meningitis caused by various pathogens: bacterial (36 publications: tuberculosis, Bacillus anthracis/Anthrax, Borrelia, Clostridium, E. coli, group B streptococcus, Listeria), parasitic (20 publications, among them Angiostrongylus cantonensis, amoeba, Cryptococcus, Toxocariasis, and Toxoplasmosis), viral (18 publications: HIV, HTLV, SARS-CoV-2, Epstein-Barr virus, Murray Valley encephalitis, tick-borne encephalitis virus, Nipah virus, respiratory viruses [various strains], West Nile virus, and Enterovirus), and fungal (14 publications: Candida, Coccidioides, Blastomyces, and Histoplasma).

##### 4.1.2. LME pattern

32 studies did not report on the LME pattern, while 4 studies did (2 spread, 2 nodular). 33 studies did not report on the LME evolution over follow-up, while 3 studies reported LME increase with clinical worsening, and 1 study in cryptococcal meningitis reported LME resolution with clinical improvement.(37)

##### 4.1.3. Imaging

Most studies employed a postcontrast T1w sequence to visualize LME (16 publications), followed by T2w-FLAIR (7 publications). 11 studies did not report which sequences were used for LME detection.

##### 4.1.4. Meta-analysis

A meta-analysis on LME in infectious diseases, including a total of 831 cases, showed an overall proportion of 0.59 [95%-CI: 0.47–0.69] with substantial heterogeneity across studies (I^2^=86%, p<0.01) (**Figure 6**). COVID-19 had the lowest proportion of LME with 0.24 [95%-CI 0.13–0.41].

**Figure 6:**
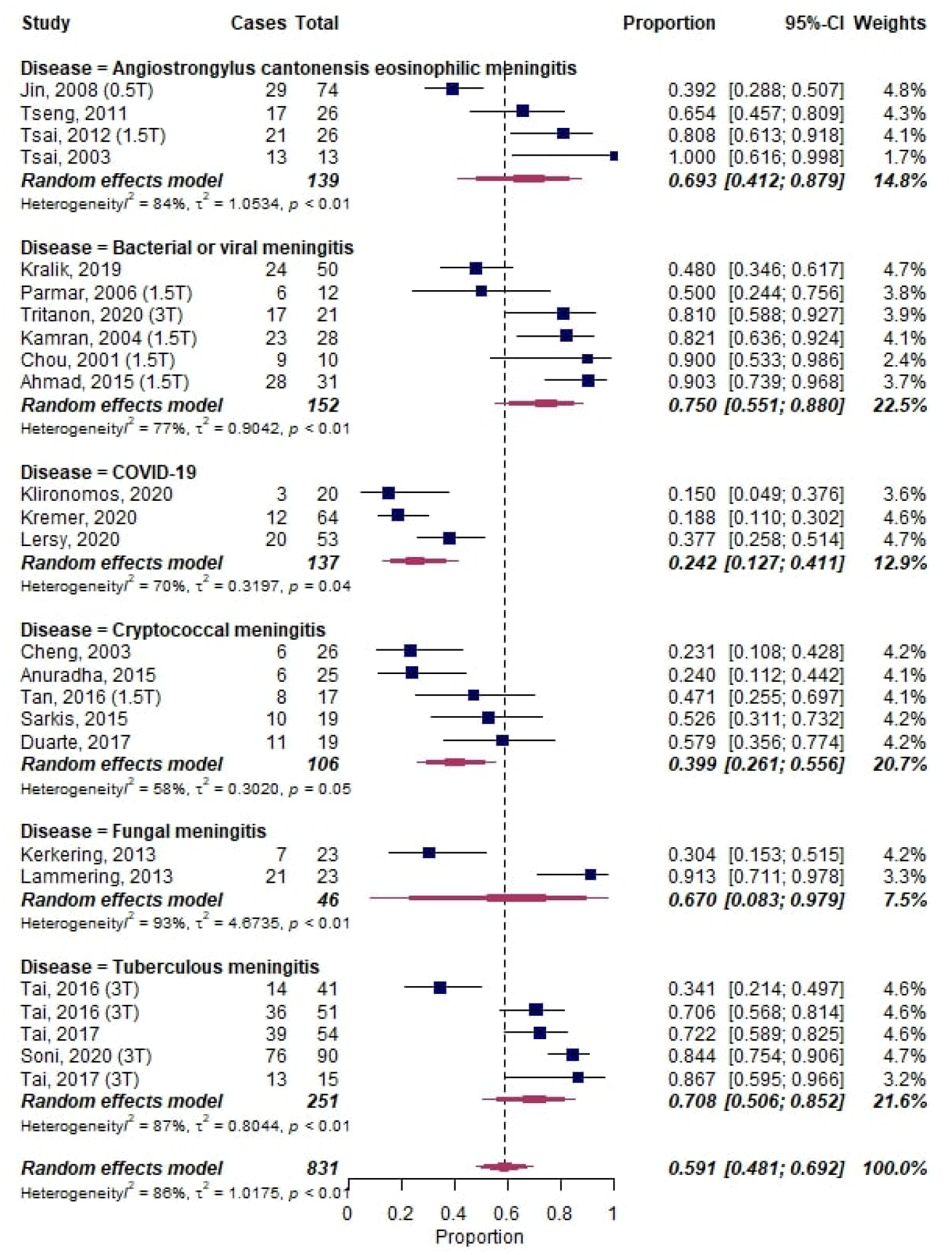
Forest plot of leptomeningeal enhancement (LME) proportions in infectious neurological diseases. Pooled analyses of studies comparing the proportion of LME on MRI in infectious neurological diseases. Static magnetic field strength for MRI acquisition for respective studies are listed in brackets if reported. Proportions for LME were extracted and pooled using the random effects DerSimonian-Laird method. Abbreviations: CI confidence interval.

#### 4.2. Neoplastic CNS diseases

##### 4.2.1. Diseases

Studies reporting on LME in neoplastic CNS diseases were caused by primary CNS tumors (92 publications: CNS lymphoma, choroid plexus papilloma, meningioma, germinoma, lipoma, primitive neuroectodermal tumors (PNET), diffuse leptomeningeal glioneuronal tumor, midline glioma, hemangioblastoma, glioblastoma/high-grade astrocytoma, Hodgkin lymphoma, xanthogranuloma, medulloblastoma, melanoma, oligodendroglioma, pilocytic astrocytoma, anaplastic astrocytoma, and giant cell astrocytoma), leptomeningeal metastases (47 publications: metastases from breast cancer, acute myeloid leukemia, rhabdomyosarcoma, gastric cancer, lung adenocarcinoma, small-cell lung cancer, pancreatic cancer, melanoma, Waldenstrom macroglobulinemia [Bing-Neel syndrome], and multiple myeloma), and hereditary tumor syndromes (5 publications: von Hippel-Lindau syndrome, Klippel-Trenaunay-Weber syndrome, and tuberous sclerosis).

##### 4.2.2. LME pattern

28 studies did not report on the LME pattern, while 8 studies did (3 spread, 3 nodular, 2 laminar/linear). 33 studies did not report on LME evolution over follow-up, while 1 study in glioblastoma reported persistent LME at follow-up MRI (up to two years later).(38)

##### 4.2.3. Imaging

Most studies employed a postcontrast T1w sequence to visualize LME (18 publications) followed by a postcontrast T2w-FLAIR (6 publications). 14 studies did not report which sequences were used for LME detection.

##### 4.2.4. Meta-analysis

A meta-analysis on LME in neoplastic diseases, including a total of 1393 cases, showed an overall proportion of 0.47 [95%-CI: 0.37–0.57] with substantial heterogeneity across studies (I^2^=90%, p<0.01) (**Figure 7**). CNS leukemia had the lowest proportion of LME with 0.24 [95%-CI 0.13–0.39]. Bing-Neel syndrome (Waldenstrom macroglobulinemia) showed the highest proportion of LME with 0.74 [95%-CI 0.60–0.84].

**Figure 7:**
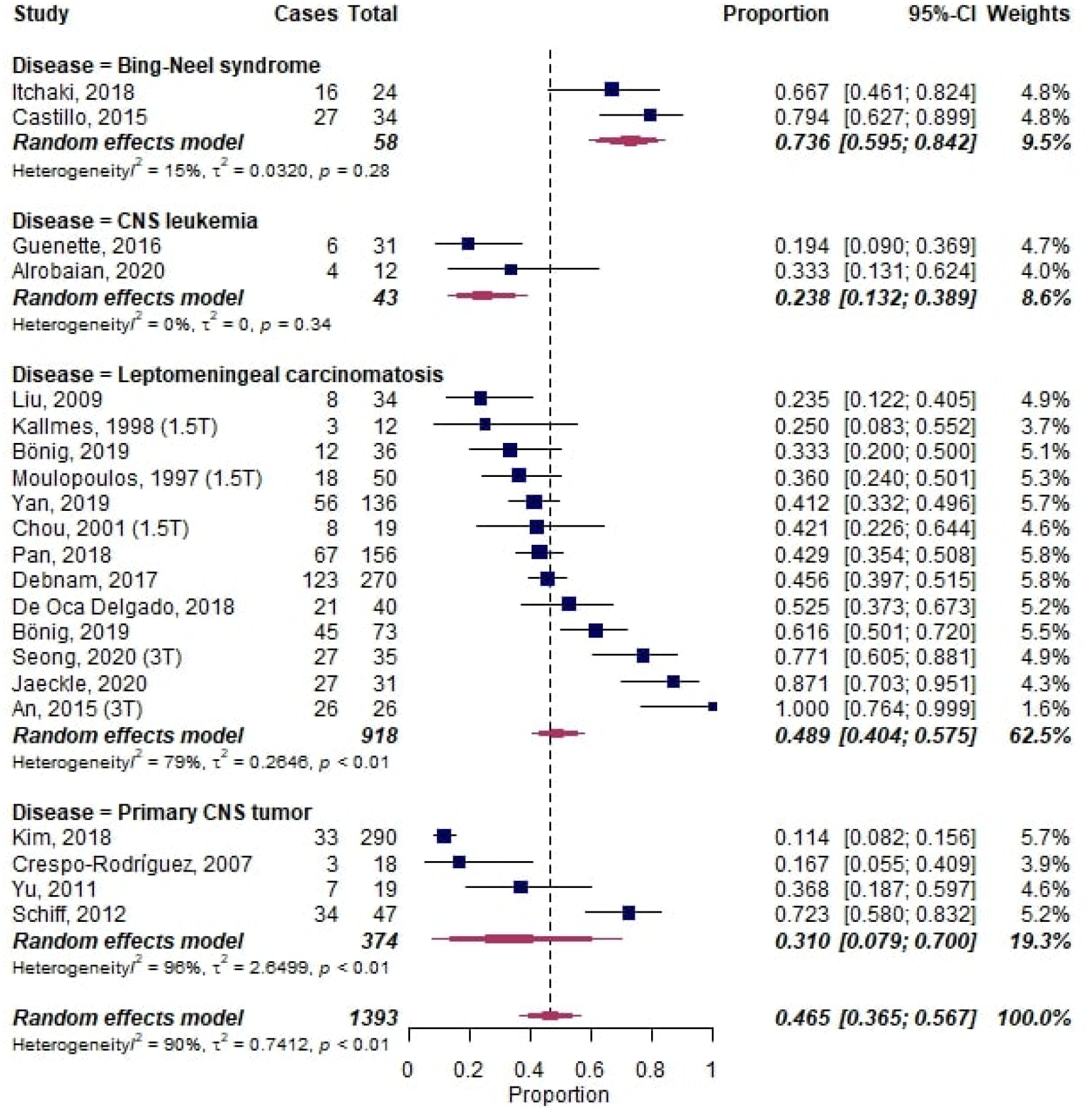
Forest plot of leptomeningeal enhancement (LME) proportions in neoplastic neurological diseases. Pooled analyses of studies comparing the proportion of LME on MRI in infectious neurological diseases. Static magnetic field strength for MRI acquisition for respective studies are listed in brackets if reported. Proportions for LME were extracted and pooled using the random effects DerSimonian-Laird method. Abbreviations: CI confidence interval; CNS, central nervous system.

#### 4.3. Other neurological diseases including vascular diseases

##### 4.3.1. Diseases

Of 147 publications, the most notable neurological diseases not belonging to the classes above were: rheumatoid arthritis with meningitis (16 publications), Sturge-Weber syndrome (16 publications), familial leptomeningeal amyloidosis/polyneuropathy (11 publications), ischemic (reversible cerebral vasoconstriction syndrome, stroke, post interventional revascularization, severe carotid stenosis, 11 publications), cerebral amyloid angiopathy (7 publications), epileptic seizures (6 publications), Moyamoya disease (6 publications), intoxications/drug-induced LME (abrin, ibuprofen, ipilimumab, propofol [in children], tacrolimus; 5 publications), hemophagocytic lymphohistiocytosis (4 publications), posterior reversible encephalopathy syndrome (PRES) (4 publications), Rosai-Dorfman disease (3 publications), hepatic encephalopathy (3 publications), Sjogren syndrome (2 publications), traumatic brain injury (2 publications).

##### 4.3.2. Meta-analysis

A meta-analysis on diseases with ≥2 publications and ≥10 subjects per study, including a total of 187 cases, showed an LME proportion of 0.31 [95%-CI 0.16–0.52] in cerebral amyloid angiopathy and 0.69 [95%-CI 0.27–0.93] in reversible cerebral vasoconstriction syndrome (**Figure 8A**).(39, 40)

**Figure 8:**
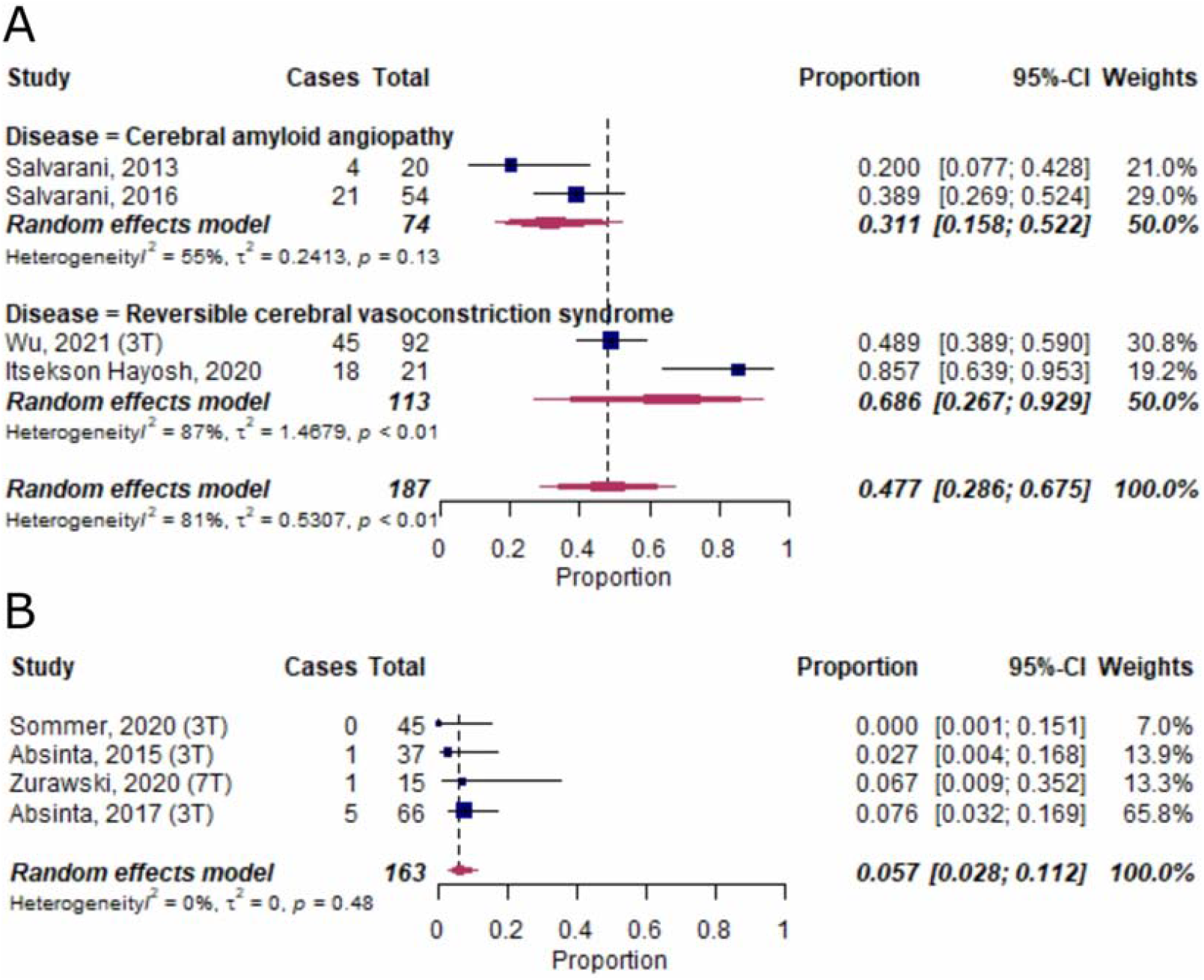
Forest plot of leptomeningeal enhancement (LME) proportions in neurological diseases not classified into the other groups and in healthy controls. Pooled analyses of studies comparing the proportion of LME on MRI in other neurological diseases (A) and in healthy control subjects (B). Static magnetic field strength for MRI acquisition for respective studies are listed in brackets if reported. Proportions for LME were extracted and pooled using the random effects DerSimonian-Laird method. Abbreviations: CI confidence interval.

### 5. Leptomeningeal enhancement in healthy controls

LME has also been reported in healthy control subjects (6 publications). A meta-analysis of 4 publications, including a total of 163 individuals, corroborated the presence of LME in this group, albeit at a low overall proportion of around 0.06 [95%-CI 0.03–0.11] (**Figure 8B**). In addition to the low proportions of LME, it has been shown in 2 publications that none of the LME-positive control subjects had more than 1 LME focus.(13, 18) In another publication, the 2 LME-positive controls exclusively presented with more than one nodular LME foci.(41)

## Discussion

### Main findings

This study aims to provide systematic evidence on LME proportion in neurological diseases, including MS, and to determine whether the role of LME as a prognostic biomarker for MS is substantiated in the literature. Overall, primary neuroinflammatory diseases showed lower LME proportion (0.26 [95%-CI: 0.20–0.35]) compared to neoplastic (0.47 [95%-CI: 0.37–0.57]) or infectious neurological diseases (0.59 [95%-CI: 0.47–0.69]). Additionally, the presence of LME was associated with worse clinical and imaging parameters in MS, that is, on average MS patients with LME had 2 years longer disease duration (p=0.03), higher EDSS by 0.7 points (p=0.006), 21 ml less cortical volume (p=0.002), 5.9 ml more T2 lesion volume (p<0.001), and 1.6 ml more T1 lesion volume (p=0.04) compared to MS patients without LME. Finally, based on a few histopathological validation studies in MS and neuroinflammatory animal models, LME corresponds to meningeal inflammatory infiltrates as well as microglial activation in the adjacent cortex. However, the evidence supporting the association of LME with cortical MS pathology remains conflicting.

### Findings in the context of existing evidence

A wide variety of disorders may present with LME, including neoplastic and infectious neurological diseases, making LME a highly nonspecific imaging finding. However, proportions of LME have considerable ranges across different neurological diseases. High LME proportions (on the order of 0.75), have been observed in Bing-Neel syndrome (a rare complication of Waldenstrom’s macroglobulinemia) (42, 43) and infectious meningitis. (44–46) Interestingly, a subset of smaller studies employing ultrahigh-field (7T) static magnetic field strengths also found proportions of LME around 0.8 in MS,(13, 16, 18) which may indicate a need for higher static magnetic field strengths to facilitate LME detection.

Other notable diseases with LME include ischemic neurological diseases, such as reversible cerebral vasoconstriction syndrome (proportion around 0.7) (39, 40) and stroke (not included in the meta-analysis).(47–49) The presence of LME in brain ischemia also suggests a relevant role of the leptomeningeal compartment in its pathogenesis. Here, LME on post-contrast T2w-FLAIR has been attributed to early blood-brain-barrier disruption,(47) also being associated with hemorrhagic transformation and worse clinical outcomes.(47) Furthermore, poor leptomeningeal collateral flow has been associated with worse clinical outcome in acute stroke.(50) Finally, also COVID-19 has been reported to present with LME, albeit with low proportions around 0.25.(51–53)

Several primary neuroinflammatory diseases can also present with LME, among them neurosarcoidosis (proportion around 0.4), MS (0.3), primary angiitis of the CNS (0.2), NMOSD (0.06), and Susac’s syndrome (not included in the meta-analysis).(54) Of note, in MS, LME proportions seem to vary among clinical phenotypes, with relapsing-remitting having lower proportions than progressive MS (0.2 vs. 0.4). However, overall longer disease duration in progressive MS could be a confounding factor.

Our meta-analysis substantiates the role of LME as prognostic biomarker in MS. The presence of LME was associated with worse physical disability and higher lesion burden as well as lower cortical volumes — the latter being also associated with worse clinical MS outcomes (reviewed in (55)). With this, our study highlights the relevance of including LME in routine clinical imaging. Along these lines, ultrahigh-field imaging at 7T might substantially improve the sensitivity to detect LME in MS in the clinical setting, as shown by our meta-analysis.

One major remaining question about LME is its underlying tissue signature. In neoplastic and infectious CNS diseases, LME likely corresponds to increased local blood supply and/or extravasation of gadolinium. However, in neuroinflammatory diseases, the pathological substrate of LME is much less clear. Based on limited EAE and MS histopathology data, LME corresponds to meningeal inflammatory infiltrates and/or tertiary lymphoid follicles (reviewed in (1)).(30, 36) Despite conflicting evidence as to whether LME is spatially associated with cortical pathology in MS,(5) there has been a consistent association of LME with low cortical volumes across studies, also substantiated by our metaanalysis. This indicates that the pathology underlying LME could exert a diffusely deleterious effect on cortical gray matter.

Different LME patterns have been described in MS (41) and, to a much lesser extent, in other neuroinflammatory diseases such as Susac syndrome,(20) neurosarcoidosis,(21, 22) and NMOSD.(23–25). LME patterns were very rarely reported in non-inflammatory neurological diseases, mostly as diffuse LME in neoplastic neurological diseases.(56) In MS, the prevalence of different LME patterns, their nomenclature as well as their association to clinical measures were highly inconsistent among studies. Nevertheless, different LME patterns could represent distinct pathophysiological features, also emphasized by the observation that healthy controls may present with nodular but not non-nodular LME.(41) With this, more data and a more stringent nomenclature is needed to describe LME phenotypes and their potential association to clinical disability and disease phenotypes. Regarding nomenclature, we favor the convention of nodular versus linear LME.

It is interesting that LME has also been observed in healthy controls, albeit at low proportions (0.06).(18, 30, 57, 58) The etiology of LME foci in healthy subjects is still a matter of debate. However, one potential cause could be minor traumatic brain injuries. It should be emphasized that LME foci can be very subtle and can easily be misinterpreted on MRI. Hence, several imaging pitfalls for LME should be taken into account, among them: gadolinium leakage of indeterminate biological significance, enhancement related to slow blood flow of cortical veins, or anatomic structures as well as imaging artifacts.(15)

Finally, data from clinical studies do not suggest a therapeutic effect of DMTs on LME in MS. However, most of these studies have a small sample size and might thus be insufficiently powered to detect a potential therapeutic effect. In addition, newer DMTs such as Bruton tyrosine kinase inhibitors (59) have not been assessed in this regard, even though these drugs led to a resolution of LME in one rodent study employing a neuroinflammatory model.(36). Hence, more data is needed on potential therapeutic impact of DMT on LME resolution. However — in our opinion — LME should still be routinely assessed in MS patients in order to enhance the knowledge and experience of radiologists and referring neurologists on this matter.(60)

### Limitations

Our study has some limitations: First, a wide variety of imaging methods have been employed to detect LME, for which we only partially corrected our analysis (e.g., static magnetic field strengths). Notably, the use of either T1w or T2w-FLAIR postcontrast sequences was not considered, with the latter generally having higher sensitivity to detect LME.(4, 19) This could have led to an underestimation of LME proportions in studies acquiring T1w sequences. However, studies in neoplastic and infectious neurological diseases with mostly bulk LME mainly employed T1w sequences, potentially counterbalancing this effect. Along these lines, we also point out that a substantial number of studies did not report on which MRI sequences they employed (T1w versus T2w-FLAIR). Second, for assessing the prognostic value of LME, we pooled studies with various methodological backgrounds for summary estimates. Nonetheless, the outcome measures reported were surprisingly uniform, even allowing for the use of mean differences in our meta-analysis.

### Conclusions

Our study provides systematic evidence for LME proportions in a comprehensive panel of neurological diseases, including MS. This high-level evidence also corroborates the prognistic value of LME in MS, supporting the inclusion of LME as a standard imaging feature in clinical MS imaging. Furthermore, this systematic review indicates that future LME studies will need to control for static magnetic field strength, type of 3D T2w-FLAIR sequence, scanner type, timing of acquisition, prior steroid administration as well as exclusion of LME imaging mimicks. More evidence on the pathological substrate of LME and on its potential association with cortical pathology in neuroinflammation is needed to further improve our understanding for this MRI feature and to strengthen its role in the clinical and research settings.

## Acknowledgements

We thank Irene Cortese, Govind Nair, Avindra Nath, and Bryan Smith for facilitating MRI acquisition. We also thank the Intramural Research Program of National Institute of Neurological Disorders and Stroke for financial support.

## Glossary

CIS: clinically isolated syndrome
CNS: central nervous system
EAE: experimental autoimmune encephalomyelitis
EDSS: Expanded Disability Status Scale
FLAIR: fluid-attenuated inversion recovery
Gd: gadolinium
LME: Leptomeningeal enhancement
MRI: magnetic resonance imaging
MS: multiple sclerosis
(RR: relapsing-remitting)
NMOSD: neuromyelitis optica spectrum disorder

## Compliance with Ethical Standards

### Data sharing statement

Data available from the authors upon request.

### Code availability

R code for meta-analysis available upon request.

## Supplementary figure

**Figure S1:**
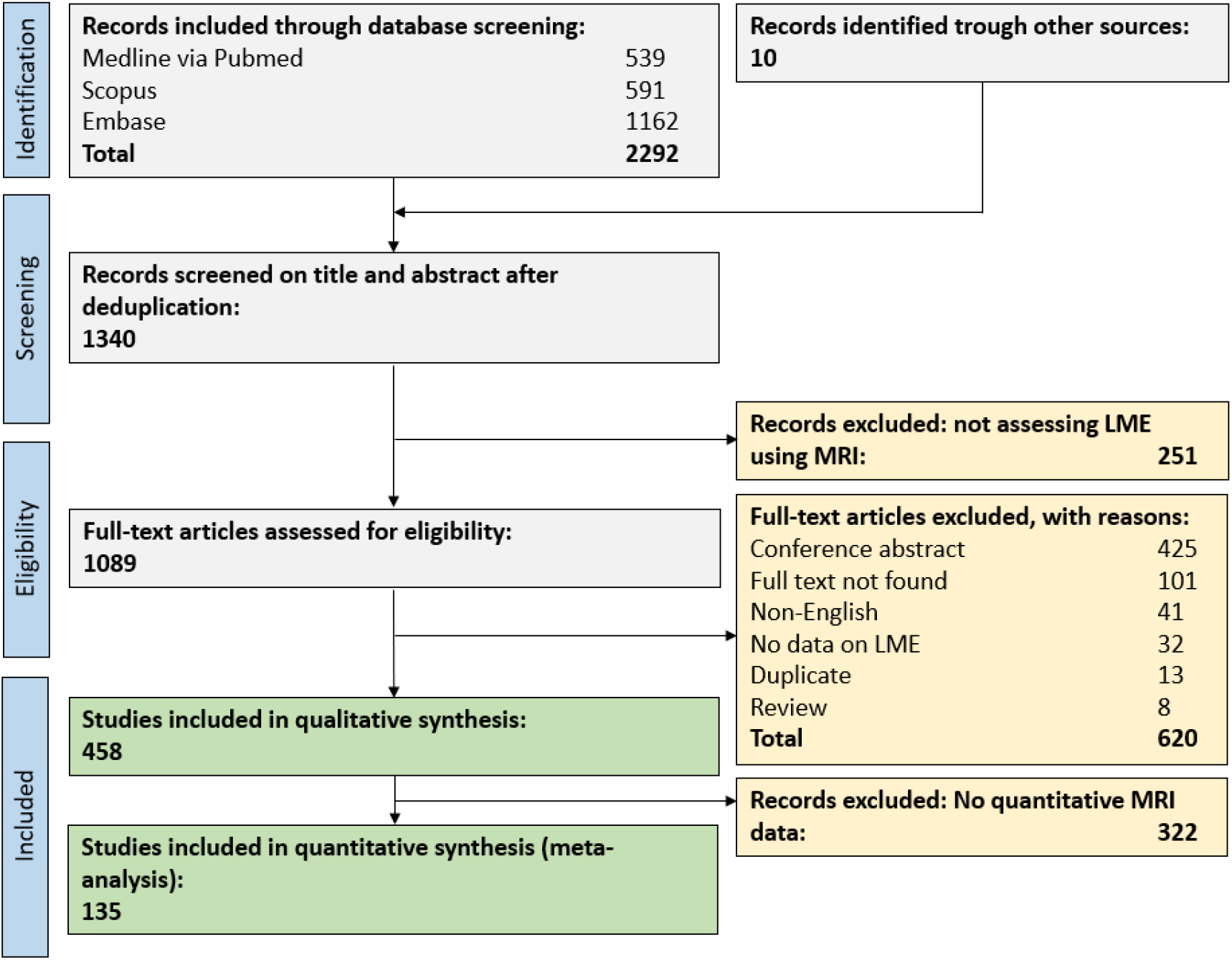
Flow chart for study inclusion.

## Supplementary data

### Supplementary search string

In alphabetical order

## Supplementary tables

**Table S1:** Predefined systematic search strings in PubMed Ovid EMBASE and Web of Science. Last search 2^nd^ of February, 2021.

| Databases | Studies produced | Search string |
| --- | --- | --- |
| <u>Pubmed via Medline:</u> | 539 | Leptomeningeal Enhancement OR Leptomeningeal Contrast Enhancement OR Leptomeningeal Inflammation OR pericortical enhancement |
| <u>Scopus:</u> | 591 | Leptomeningeal Enhancement OR Leptomeningeal Contrast Enhancement OR Leptomeningeal Inflammation OR pericortical enhancement |
| <u>Web of Science:</u> | 1162 | Leptomeningeal Enhancement OR Leptomeningeal Contrast Enhancement OR Leptomeningeal Inflammation OR pericortical enhancement |

**Table S1:**
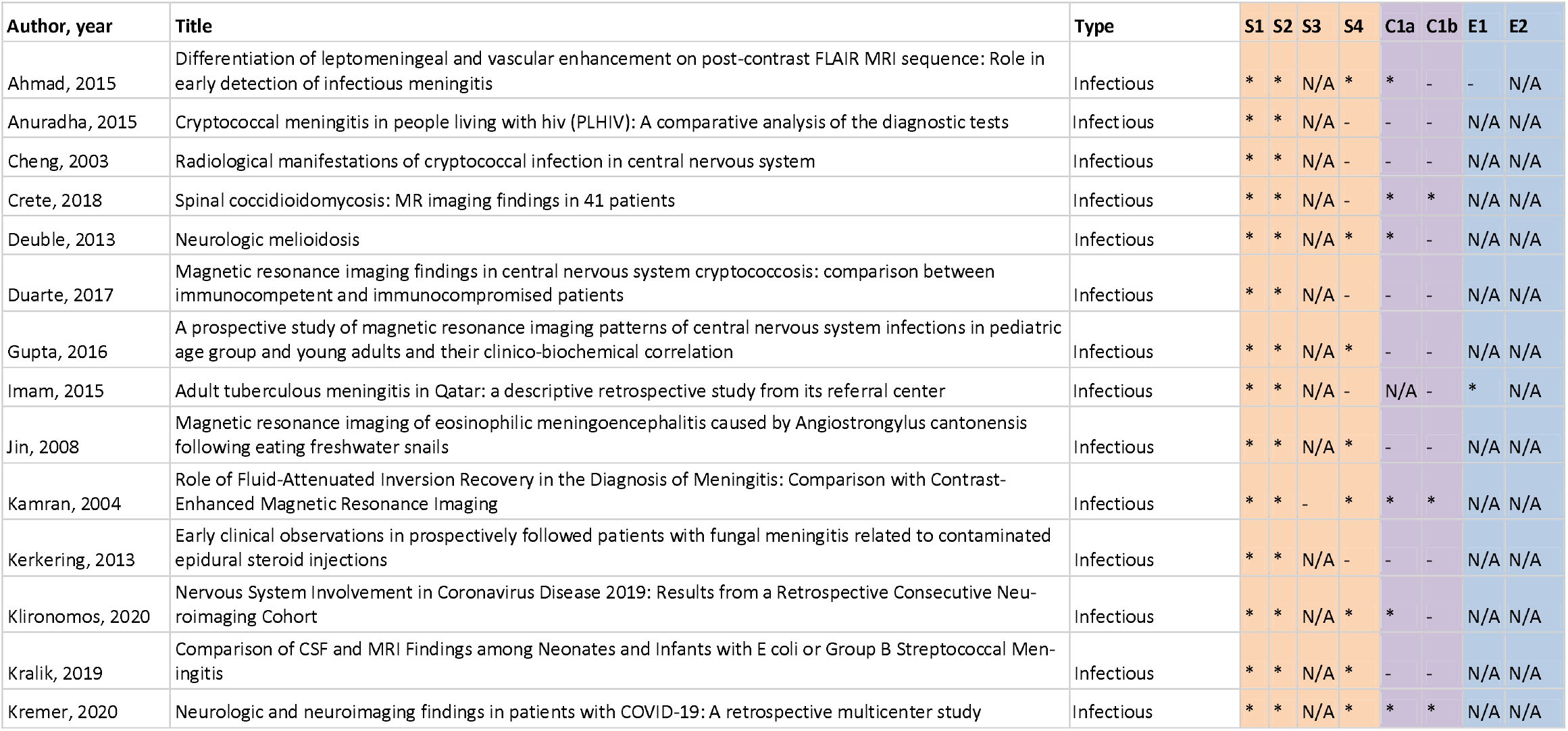
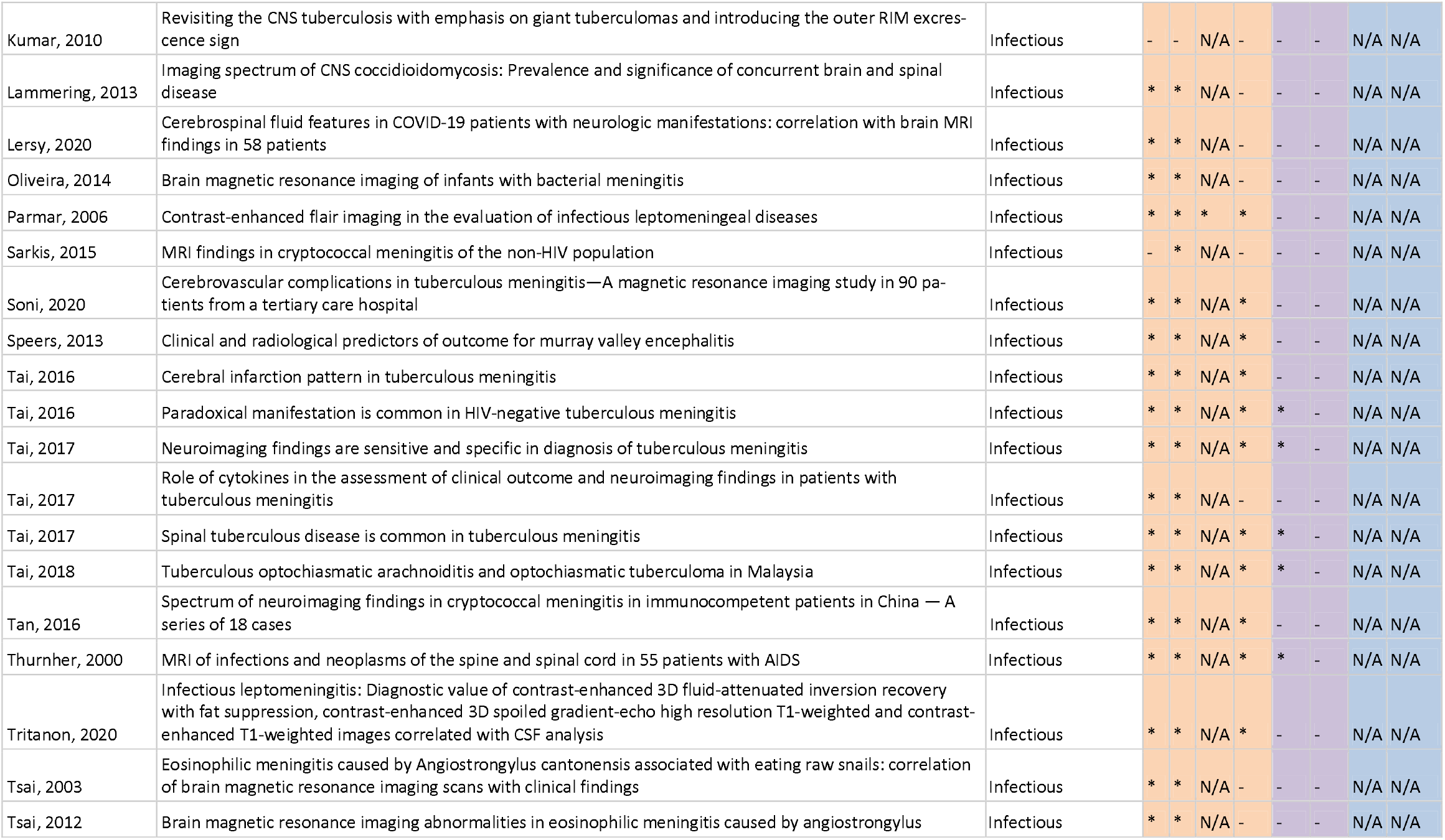
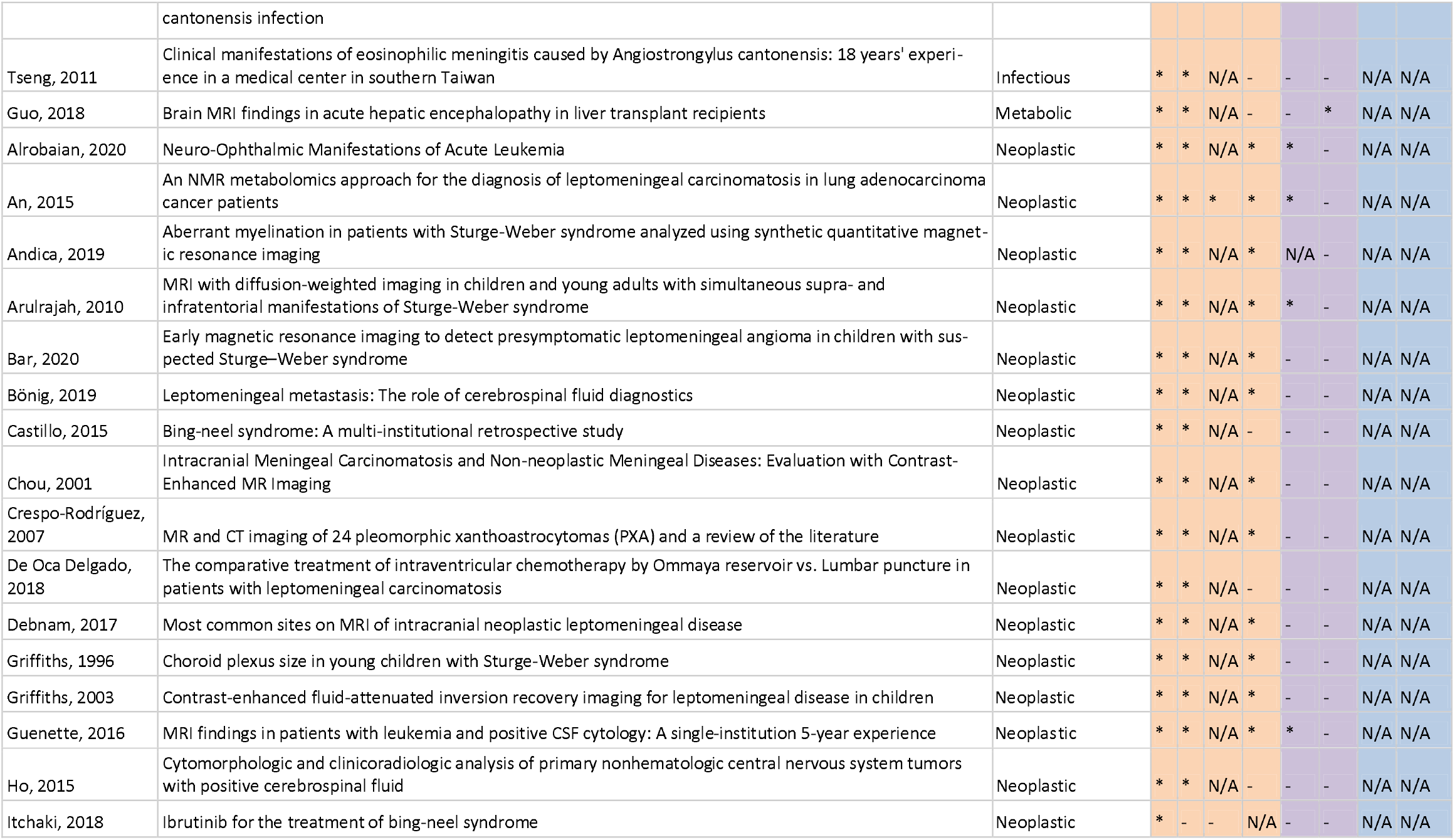
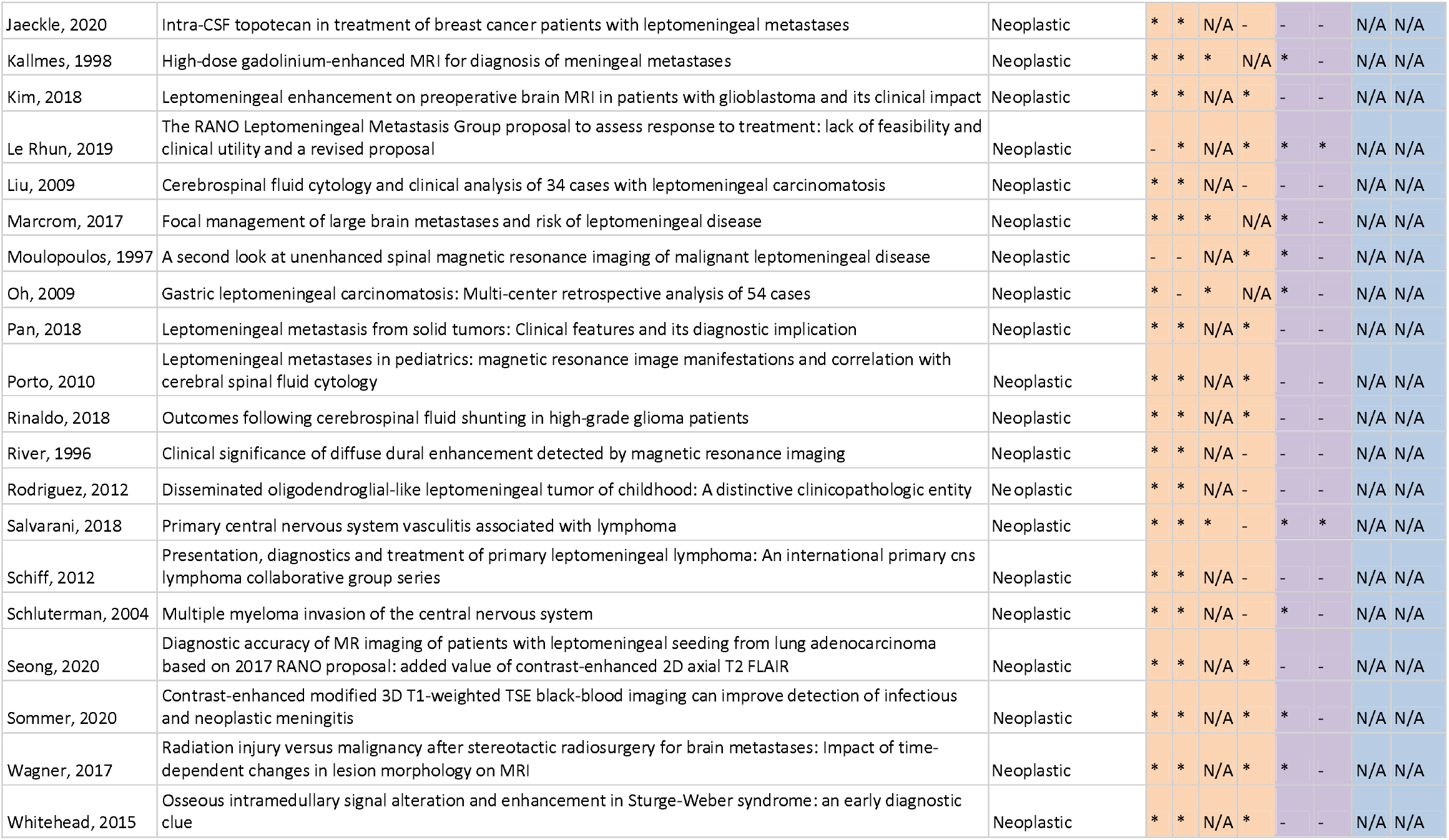
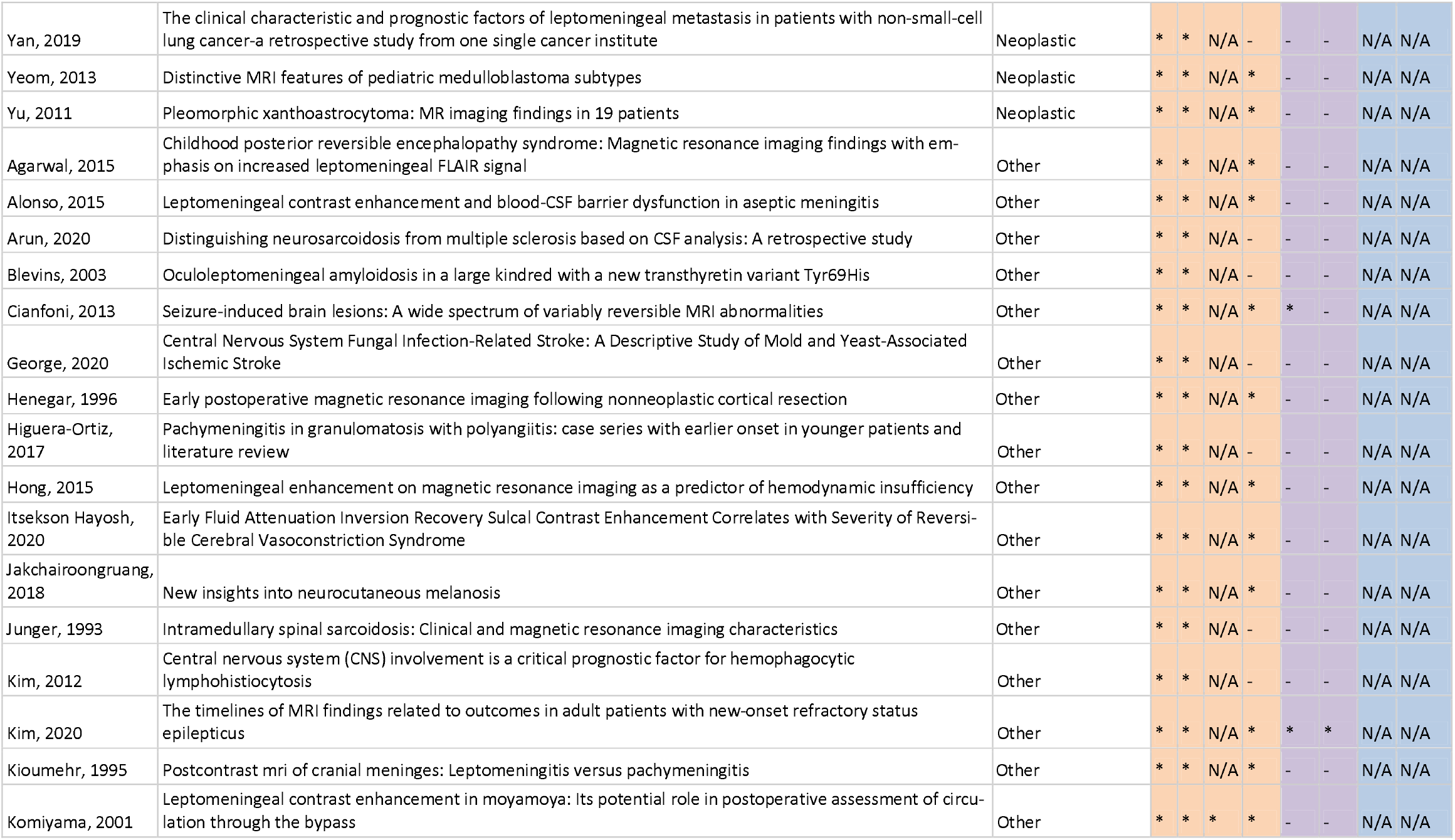
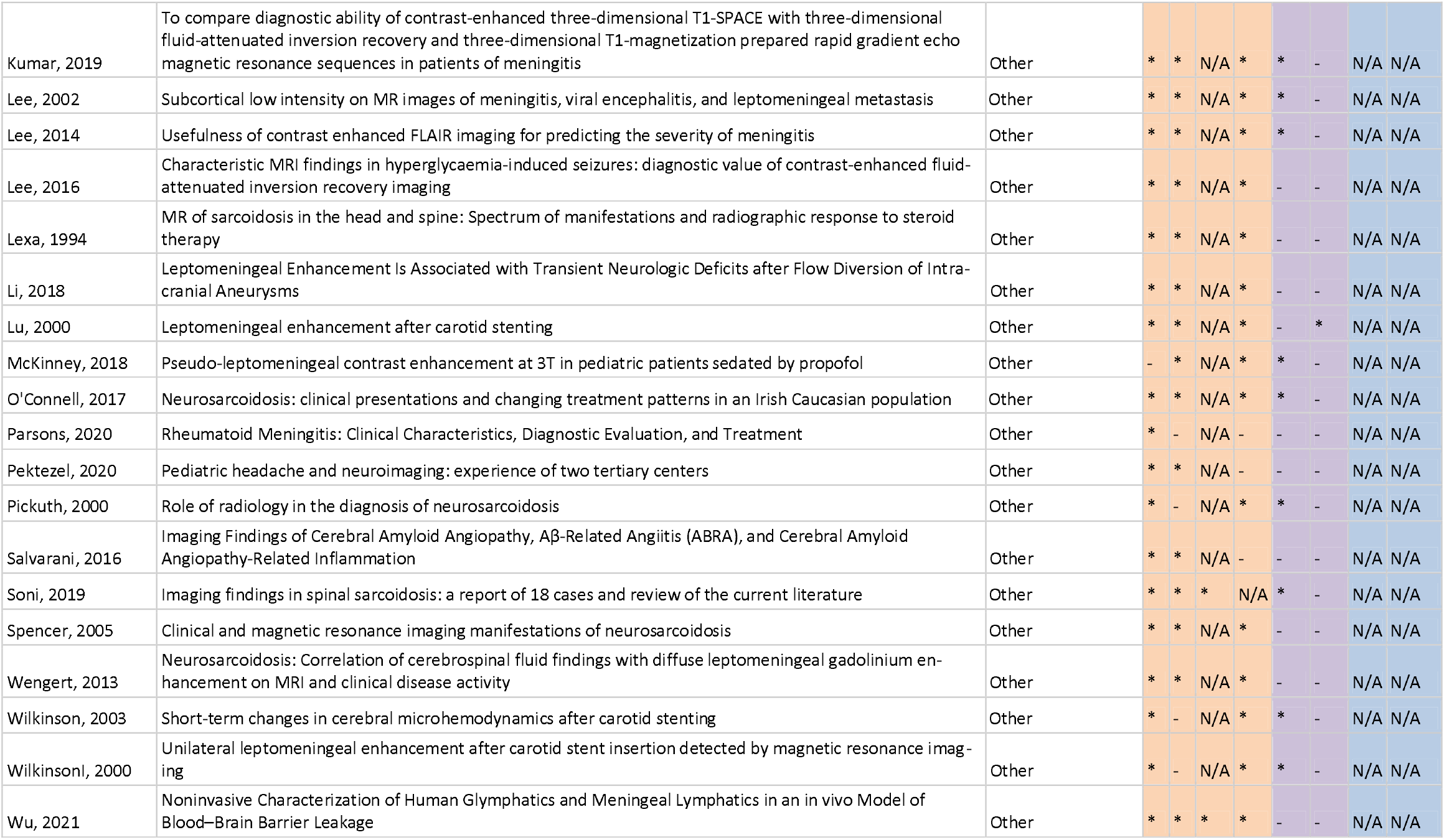
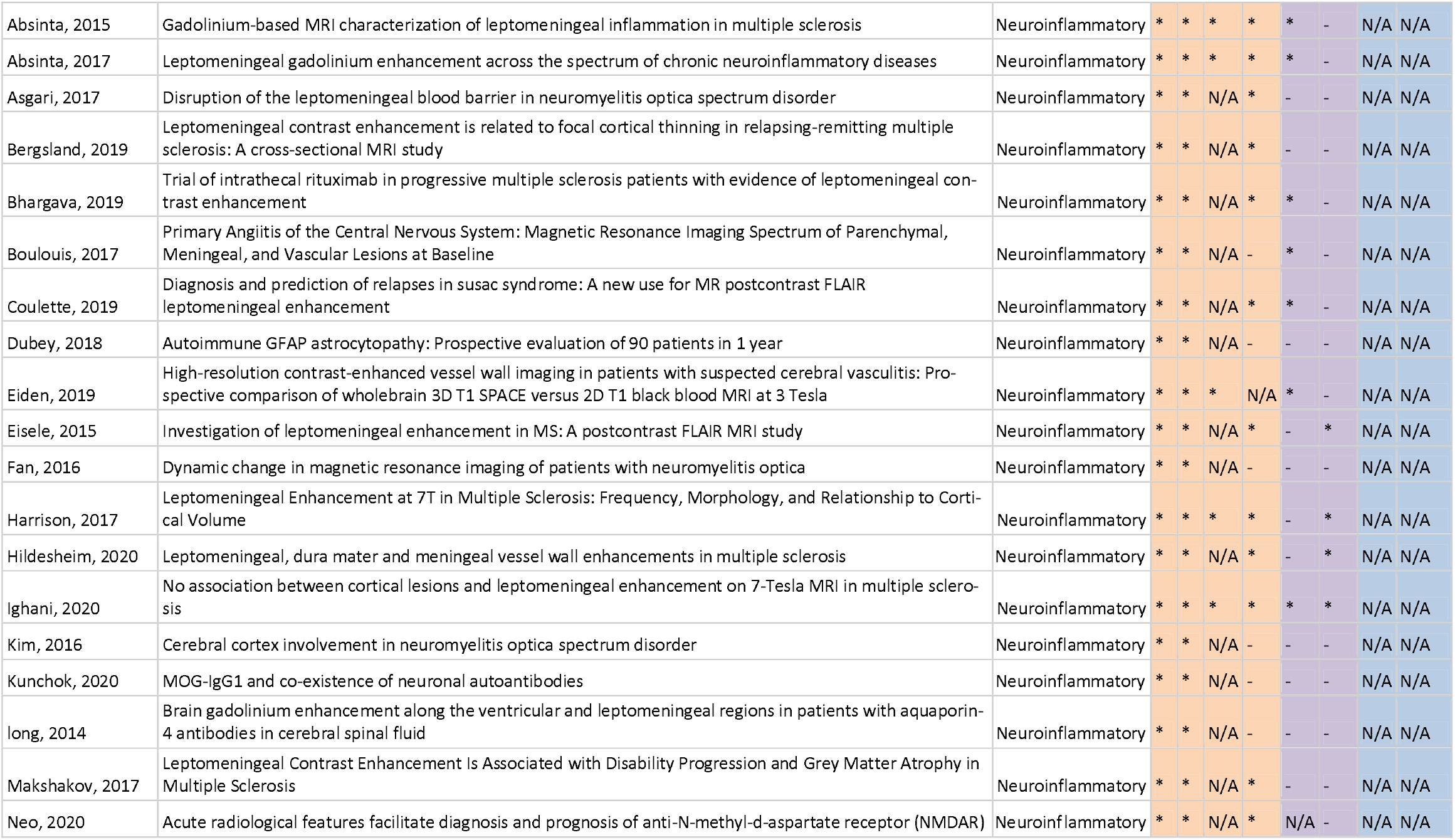
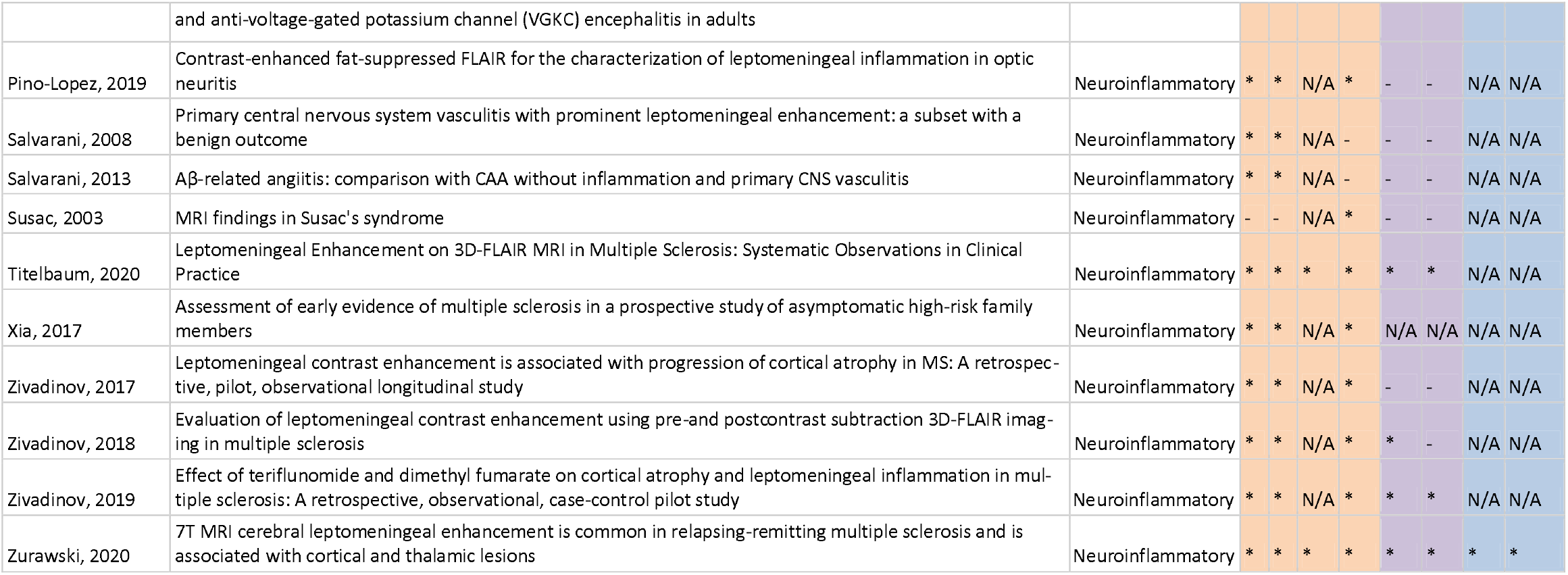
Risk of bias assessment of the included studies with 10 or more included human subjects according to the Newcastle-Ottawa sea le for nonrandomized studies. S1-S4: Selection domain (only S1-S3 for cohort studies, case series and retrospective studies), C1a and C1b: comparability domain, E1-E2: exposure domain. In alphabetical order and sorted for disease class.

**Table S3:**
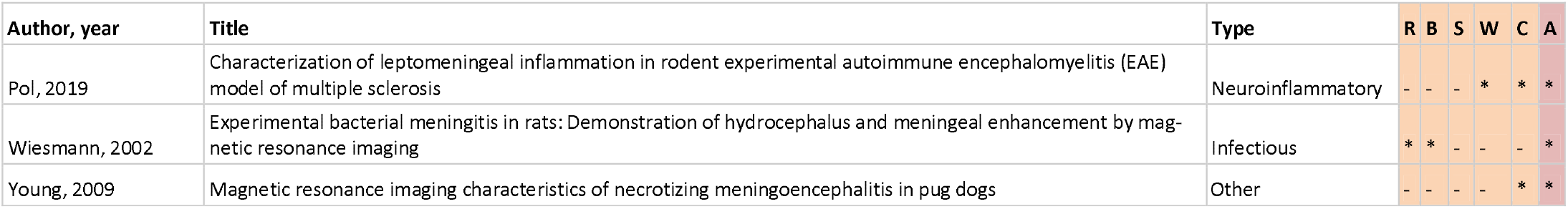
Risk of bias assessment of the included animal studies according to the risk of bias assessment checklist for good research practice and whether a study was in accordance with the ARRIVE guidlines. A, in accordance with ARRIVE guidelines; B, blinding; C, conflict of interest; R, randomization; S, prior sample size calculation; W, animal welfare statement.

