## Supplementary figures and images for "Leptomeningeal Enhancement in Multiple Sclerosis and Other Neurological Diseases: A Systematic Review and Meta-Analysis"

### Supplementary figure 1

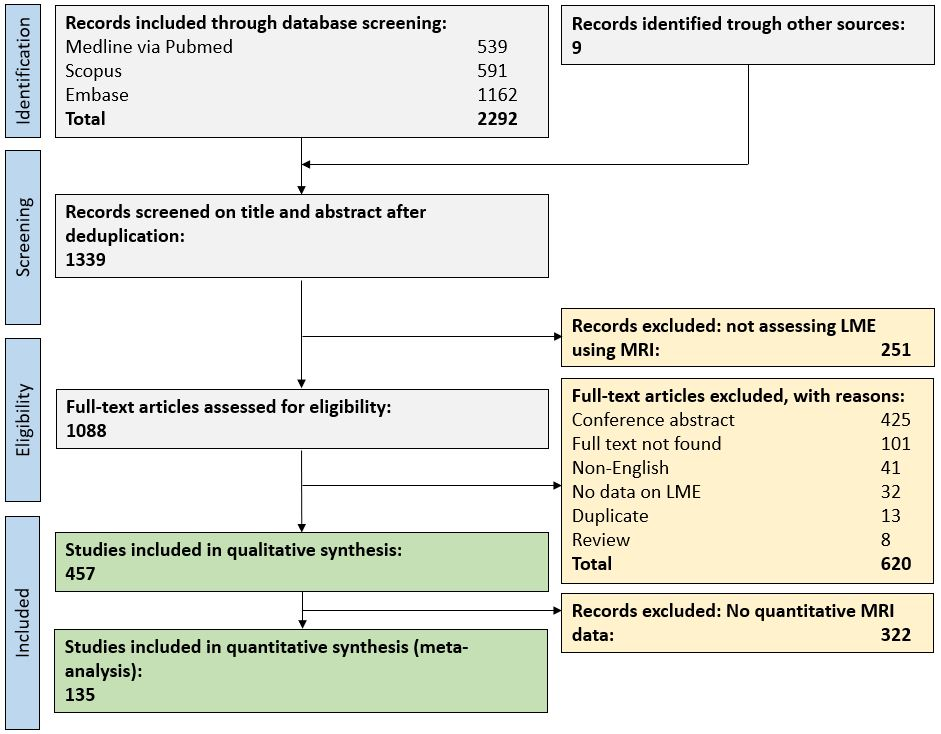
